# IgG-inspired, multivalent protein-DNA nanostructures for high-affinity, tunable, and reversible binding to biomolecular targets

**DOI:** 10.1101/2023.09.18.558353

**Authors:** Rong Zheng, Yang Xu, Abhay Prasad, Minghui Liu, Zijian Wan, Jiapei Jiang, Xiaoyan Zhou, Ryan M. Porter, Matthew Sample, Erik Poppleton, Jonah Procyk, Hao Liu, Adam Doherty, Pranish Nyaupane, Yize Li, Shaopeng Wang, Hao Yan, Petr Šulc, Nicholas Stephanopoulos

**Affiliations:** School of Molecular Sciences, Center for Molecular Design and Biomimetics, The Biodesign Institute, Arizona State University, Tempe, AZ 85281, USA; School of Biological and Health Systems Engineering, Center for Biosensors and Bioelectronics, The Biodesign Institute, Arizona State University, Tempe, Arizona 85281, USA; School of Electrical, Computer and Energy Engineering, Biodesign Center for Biosensors and Bioelectronics, Arizona State University, Tempe, Arizona 85287, USA; School of Life Sciences, ASU-Banner Neurodegenerative Disease Research Center, The Biodesign Institute, Arizona State University, Tempe, AZ-85281, USA; John Shufeldt School of Medicine and Medical Engineering, Tempe, AZ 85287, USA; School of Natural Sciences, Department of Bioscience, Technical University Munich, 85748 Garching, Germany

## Abstract

Multivalency enables nanostructures to bind molecular targets with high affinity. Although IgG antibodies can be generated against a wide range of antigens, their shape and size cannot be tuned to match a given target. DNA nanotechnology provides an attractive approach for designing customized multivalent scaffolds due to the addressability and programmability of the nanostructure shape and size. Here, we use computational simulation to guide the design and synthesis of a DNA nanostructure-based synthetic antibody (“nano-synbody”). The nano-synbody is comprised of a three-helix bundle DNA nanostructure with three identical arms terminating in a mini-binder protein that targets the SARS-CoV-2 spike protein. The structure was designed to match the valence and distance between the three receptor binding domains (RBDs) in the spike trimer, in order to enhance binding through avidity effects. Moreover, the design allowed for the display of one, two, or three protein-displaying arms, thereby systematically probing the effect of multivalency on binding affinity. The binding strength of the nano-synbody increased with the increasing number of arms, yielding 11.2 pM affinity (∼100-fold enhancement over monovalent binding) for the wild-type spike protein for the three-arm structure. Moreover, the multivalency was able to yield a 95 pM affinity for the Omicron variant, a mutant against which the monovalent protein was ineffective. The nano-synbody could also block infection of a spike protein-bearing pseudovirus, and similarly demonstrated effective inhibition of the Omicron variant when trimerized. The structure of the three-arm nano-synbody bound to the Omicron variant spike trimer was solved by negative-stain transmission electron microscopy reconstruction, and shows the protein-DNA nanostructure with all three arms bound to the RBD domains, confirming the intended trivalent attachment. Finally, nano-synbody binding could be reversed by removing one, two, or three arms in a programmable fashion, via toehold-mediated strand displacement. The ability to tune the size and shape of the nano-synbody, as well as its potential ability to attach (and then remove) two or more different binding ligands, will enable the high-affinity binding of a range of proteins, and pave the way towards their manipulation using DNA-based nano-robotic devices.

## INTRODUCTION

The ability of molecules like antibodies to bind a target in a high-affinity, high-specificity manner is a cornerstone of biology and nanotechnology.^1–3^ The most commonly used antibody type for biomolecular applications—such as sensing, targeting, or imaging—is the immunoglobulin G (IgG) scaffold, consisting of the canonical “Y” shape with two identical arms, each consisting of one constant (Fc) and one variable (Fv) domain (**Figure 1A**). The presence of two arms primarily enhances binding by increasing the effective concentration of the two variable (Fv) domains, resulting in avidity through statistical rebinding.^4^ Unless the spacing between two epitopes matches the inter-Fab distance (∼10-15 nm), seen only in with rare exceptions,^5^ IgGs do not usually bind to two targets simultaneously; when this is the case, it is most often to two distinct targets on a multivalent scaffold, e.g. crosslinking two different spike complexes on a virus capsid surface.^6,7^ Such monovalent target binding to any single epitope (**Figure 1B**) renders IgGs susceptible to mutagenic escape, whereby mutations in the patch of the protein targeted by the variable regions compromise binding.^8^ This is a particularly pressing concern in the context of a rapidly evolving pathogen, like the SARS-CoV-2 virus, where subsequent variants (e.g. Delta, Omicron) abrogated binding by IgG molecules generated against the wild-type (WT) virus, resulting in waning effectiveness of induced immunity and antibody-based therapies. In addition to the fixed geometry and Fv spacing, typical IgG binding cannot be tuned dynamically in a stimulus-responsive fashion. Imparting such a capability would allow for biotechnologically useful reversible antibodies that could release their target on-demand, e.g. to “turn off” target blocking and activate a protein when desired. Moreover, reversible binding could enable release of a target from a nano-structure/particle, or serve as a nanoscale “robot arm” that can grip, move, and then release proteins.

**Figure 1:**
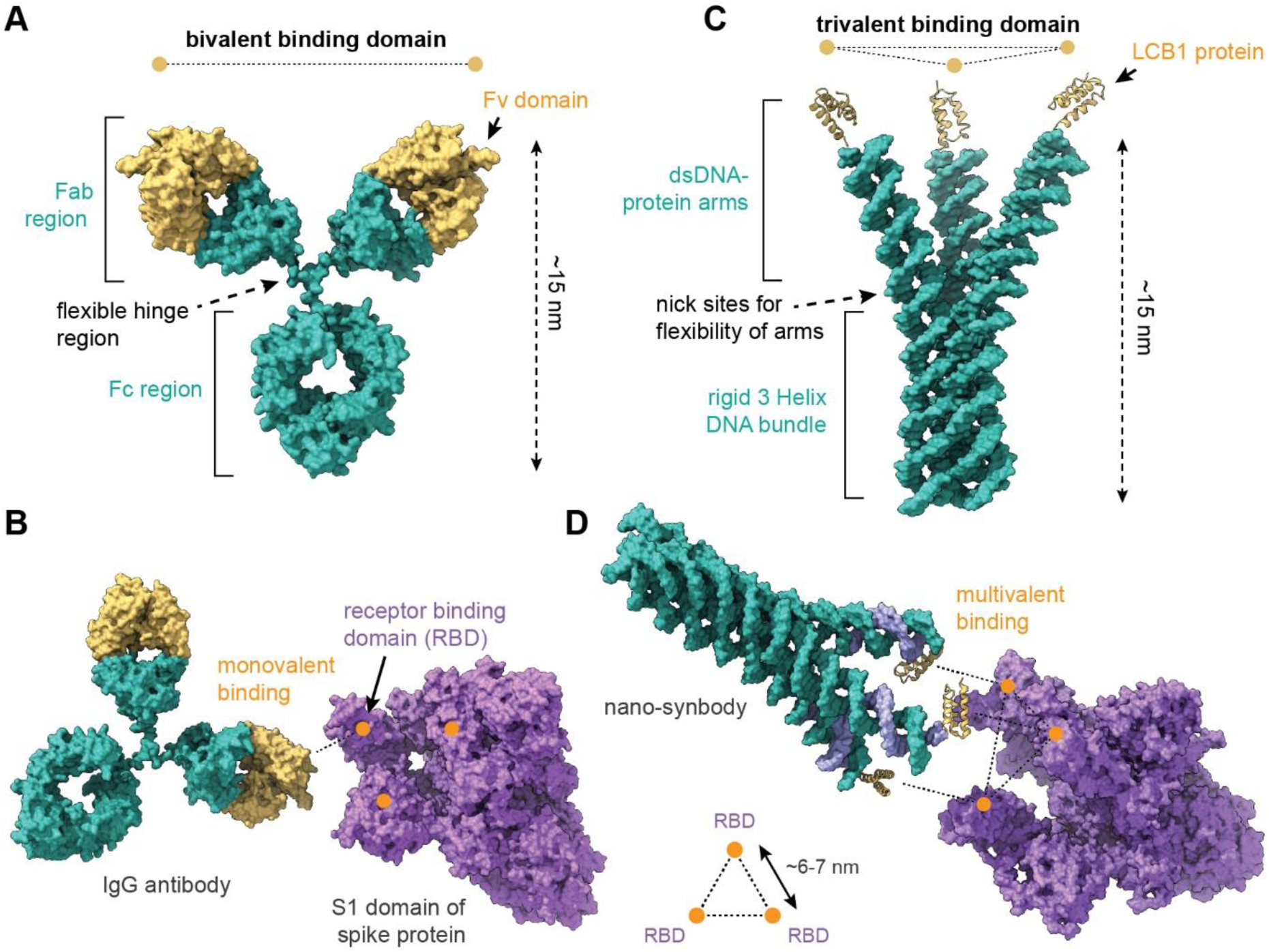
Comparison of an IgG antibody and the protein-DNA nano-synbody. **A)** Classic Y-shape of an IgG antibody, showing the bivalent presentation of two Fv domains. **B)** Structure of the three-arm nano-synbody, with three LCB1 proteins at the tips of its identical “arms.” **C)** Most IgG antibodies can only bind to a target in a monovalent fashion. **D)** The nano-synbody, by contrast, is specifically designed so that the three arms can match the distances between the three binding domains (the RBD on the SARS-CoV-2 spike trimer in this case).

We asked whether an IgG-mimetic nanostructure could be constructed that would address these two key shortcomings, enabling both multivalent association with a single biomolecular complex and reversible binding upon addition of a programmable molecular trigger. Toward this end, we turned to DNA nanotechnology, which relies on the programmable self-assembly of oligonucleotides to construct materials of precise shape, size, and functionalization.^9,10^ In the past 10-15 years, DNA linkers or nanostructures have been used extensively to control the spacing of other ligands—such as aptamers, peptides, proteins, or small molecules—and create multivalent nanostructures that can often dramatically enhance binding affinity.^11–15^ Key to this approach has been spatially matching the distribution of molecular targets with the displayed ligands, with examples ranging from 3-5 nm to tens or even hundreds of nanometers. We thus explored whether DNA nanotechnology could be used to generate an IgG mimetic nanostructure with an Fc-inspired “constant” region linked, via flexible hinges, to multiple “arms” that each bear a small Fv-mimetic binding protein.

As a proof of principle, we selected the SARS-CoV-2 spike trimer, due to the wide range of binding proteins reported for this target, and the availability of multiple spike variants with different affinity towards those proteins. Moreover, the trimeric structure of the SARS-CoV-2 spike protein presented an ideal target for nanostructured affinity agents designed to match its symmetry and spacing.^16^ Unlike IgG molecules, whose fundamental dimensions and number of Fv domains are determined by their structure, a DNA nanostructure-based synthetic antibody (with we term a “nano-synbody”, **Figure 1C**) can be tuned to match the size and valence of the target. Our design relies on creating a three-helix DNA bundle with identical arms that can hybridize to a protein-DNA conjugate, which can in turn bind to the three RBDs on the spike (**Figure 1D**). To guide the design of the nanostructure, we developed a computational pipeline—which uses a coarse-grained protein-DNA model based on the oxDNA simulation software^17–19^ —in order to ensure that the design can effectively bind the target. The nano-synbody reported here is similar in size to an IgG (∼15 nm), with inter-arm distances engineered specifically for the distance between the three RBD domains (∼7 nm).

Since the emergence of the Covid-19 pandemic, a range of DNA-based nanostructures have been reported for multivalent binding to the SARS-CoV-2 spike trimer,^20–26^ with applications ranging from sensing/diagnostics to therapeutic virus neutralizing nanostructures. These elegant examples have highlighted the power of nucleic acid scaffolds for enhancing avidity, and spatially matching the threefold symmetry of the spike complex. Our approach varies from these in that it explicitly mimics the structure of an IgG, extending the canonical “Y-shape” to three arms branching from a core constant region, and with the ability to programmably display 1-3 arms. Moreover, it is the first to demonstrate reversibility of SARS-CoV-2 binding by displacing each of the three arms in a tunable fashion. It is also, to our knowledge, the first DNA-based SARS-CoV-2 targeting nanostructure to use a mini-binder protein (as opposed to peptides, aptamers, or full-length antibodies) as the modular Fv-mimetic domain. The Fc-mimetic domain could, in turn, be used to incorporate other proteins in a modular fashion, to oligomerize the nano-synbodies for additional multivalency, or to integrate with other DNA nanostructures.

For the RBD-binding agent, we selected a *de novo* designed mini-protein (termed “LCB1”) reported in 2020 by the Baker group,^27^ and used extensively by their lab and others since.^28–31^ This choice was influenced by several factors: the compact size of the protein (7 kDa); the ease of mutagenesis and recombinant expression of the protein in *E. coli*, in order to introduce uniquely reactive residues for DNA bioconjugation; its high stability (T_m_ > 95°C); and its high-affinity binding as a monomer to the WT spike RBD, with a dissociation constant (K_D_) in the low-nanomolar range. Our results demonstrate that the binding affinity for the spike trimer increases with the number of proteins displayed, and that this binding enhancement leads to improved neutralization of a spike-bearing pseudovirus. Crucially, the multivalency afforded by the nanostructure imparts nanomolar affinity for several Omicron variants, against which the mono- or di-valent nano-synbodies (or therapeutic IgG molecules) are ineffective. To further elucidate the nano-synbody interaction with the Omicron spike (B.1.1.529), we directly imaged the structure bound to the spike trimer via negative-stain transmission electron microscopy (nsTEM). In this way, we could visualize the structure attached to the spike trimer as designed by simulation, and confirmed that all three LCB1 proteins bind as designed. Finally, we used toehold-mediated strand displacement^32^ to remove one, two, or three arms from the nano-synbody, and systematically reverse the binding of the structure to both the WT and Omicron spike variants. Taken together, our results validate our integrated computational-experimental pipeline for designing custom synthetic antibodies, and enabling tunable design of IgG-mimetic protein-DNA nanostructures against a broad range of targets in the future.

## RESULTS

### Computational design and simulation of nano-synbody structure

The SARS-CoV-2 spike trimer consists of three identical glycosylated proteins, each of which contains two functional domains, termed S1 and S2.^33^ The receptor-binding domain (RBD) located on the S1 domain plays a pivotal role in mediating the interaction between the virus and the ACE2 receptor on host cells. The conformational change of the S1 domain from the “down” to the “up” state is crucial to this process. Notably, the spatial separation between the three RBD domains in the up state is ∼7 nm, which is compatible with the distances bridged by small DNA assemblies. Our goal was to design a nanostructure with a rigid region, linked to 1-3 arms bearing the LCB1 protein, in such a way that they could effectively bind to the RBD domains. In particular, we wanted the rigid domain to be as small as possible, to minimize the number of DNA strands necessary, while still readily spanning the distances between the binding sites. Toward this end, we created a computational design pipeline outlined in **Figure 2A**. We imported the known conformational states of a target protein, which can be obtained from a crystal or cryo-EM structure, an AlphaFold prediction, or from atomistic simulation. Then, using our coarse-grained modeling, we are able to screen *in silico* a library of DNA nanostructure designs, e.g. multi-helical bundles in **Figure 2A** in order to better mimic the IgG structure. Following the determination of a multi-helical bundle core with high rigidity, structures with varying Fab-mimetic arm lengths and Fc-mimetic bundle lengths are simulated to refine our architecture into a minimal structure with optimized rigidity and arm-arm distance distributions. Through this computational design pipeline, we are able to determine the simplest nanostructure that can readily span the three binding sites, without incurring too much strain or distortion.

**Figure 2:**
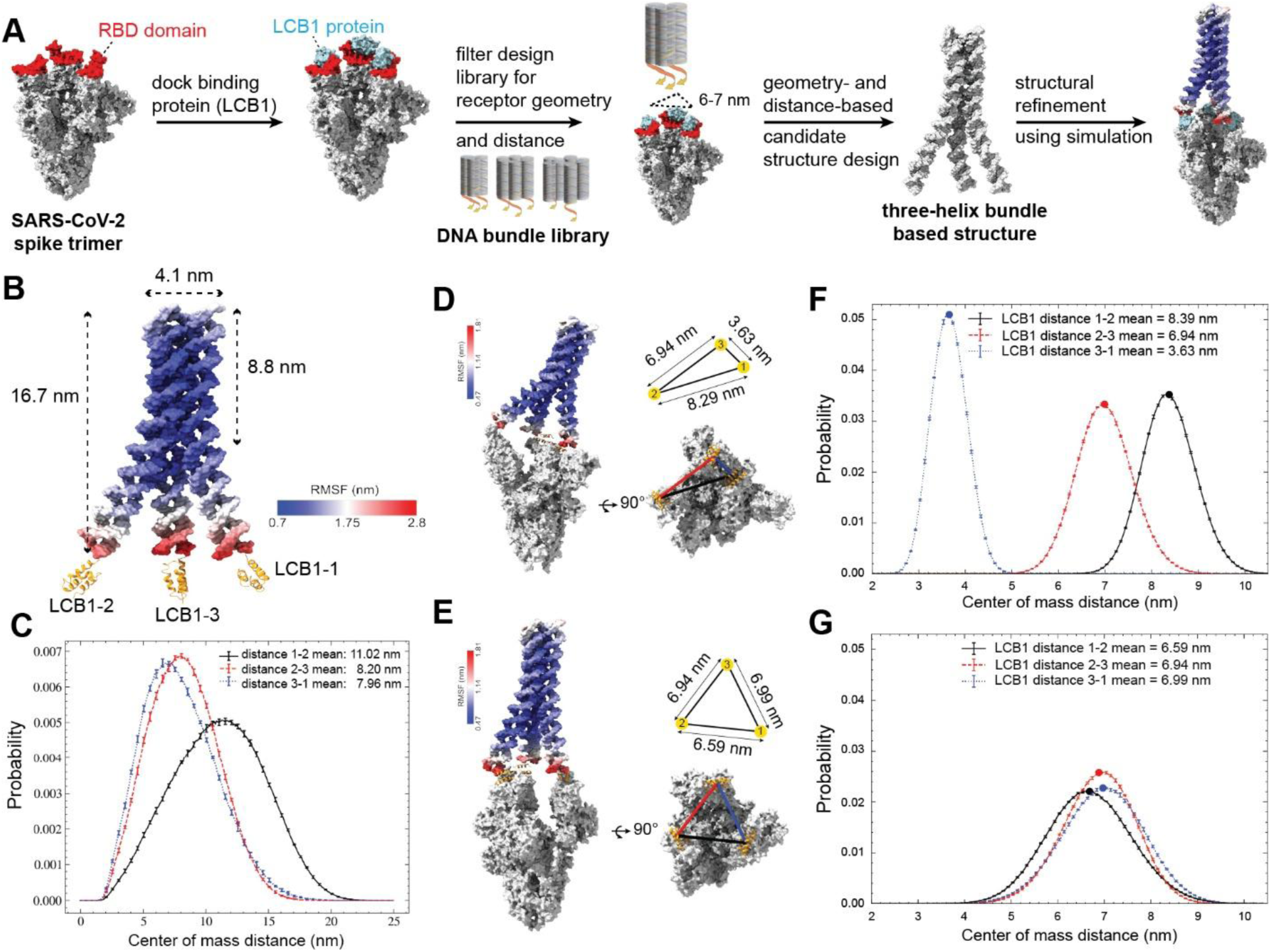
Computational simulation of the 3-arm nano-synbody. **A)** Computational design pipeline for generating the minimal nanostructure that can bridge three target sites and present the mini-binder protein LCB1. **B)** OxDNA-ANM simulation of the free 3-arm nano-synbody; coloring indicates the root mean-squared fluctuations (RMSF). **C)** Distance distributions of the three centers of mass of the LCB1 proteins, as indicated in (**B**). **D,E)** The structure of the nano-synbody bound to the spike protein in two states: with two receptor binding domains (RBDs) up and one down (**D**), and all three RBDs up (**E**). The histograms in (**F**) and (**G**) represent the distance between the center of mass of the LCB1 proteins relative to each other when bound to the spike protein in the two RBD up and three RBD up conformations, respectively. The non-overlapping distributions in (**F**) and overlapping distance distributions in (**G**) quantitatively demonstrate the nano-synbody’s capacity to adapt to different distance conditions required to enforce trivalent binding to the spike protein.

This design process suggested that a three-helix bundle (3HB)-based Fc-mimetic domain—approximately 4 nm in diameter, and fortified with several crossovers between helices to ensure rigidity—with three identical single-stranded arms was the smallest design that could display three copies of a complementary DNA-LCB1 conjugate with approximately these distances (see **Figure S1-3** for the sequences and detailed design of the 3HB structure, and **Figure S5** for comparison with other candidate nanostructures). To validate that the nano-synbody could indeed display the three proteins at the intended distance, we employed the coarse-grained model ANM-oxDNA^18^ to simulate the assembly. This model represents each nucleotide as a single rigid body, with interactions parametrized to reproduce the structural, thermodynamic and mechanical properties of single-stranded and double-stranded DNA. The model has previously been instrumental in studying a wide range of DNA nanodevices, demonstrating good agreement with experimental results where available.^17,19,34^ In the ANM-oxDNA simulation package, the dynamics of proteins are modeled using an anisotropic network model (ANM). Each amino acid is represented as a single bead, with harmonic potentials between the beads parametrized to reproduce the protein’s measured B-factors, thereby capturing its inherent flexibility. The ANM-oxDNA model has been previously shown to correctly reproduce structures of protein-DNA hybrid nanostructures, including using asymmetric harmonic potentials to model the chemical linker connecting them.^35^ Here, we used multiscale modeling combining both coarse-grained and fully atomistic simulations, to parameterize the chemical crosslinks between the nucleotides on the DNA handle and the LCB1 protein used in the experiment, and to computationally derive B-factor data of the spike proteins (see Supporting Information). We previously extended our graphical user interface tool oxView^36^ to facilitate the design and setup of protein-DNA hybrid simulations. The designed structure was then characterized *in silico* through molecular dynamics simulations to quantify its flexibility and confirm that the positions of all three LCB1 proteins were displayed suitably for binding the RBD domains on the spike trimer.

We probed both the free (i.e. unbound) nano-synbody, as well as the nano-synbody bound to the spike trimer. The simulations of the unbound nano-synbody (**Figure 2B**) showed the average width and length of the 3 helix-bundle (the Fc-mimetic domain) to be 4.1 nm x 8.8 nm respectively, while the arms (Fab-mimetic domains) had a length of ∼8.0 nm. The Fc-mimetic domain was relatively rigid, as expected from the multiple crossovers holding the three helices together. Distance distributions between the centers of mass of the three LCB1 mini-binders attached to the DNA arms (**Figure 2C**) showed a slightly asymmetric structure, with two inter-arm distances ∼8 nm, and one (between LCB1-1 and LCB1-2, in **Figure 2B**) at ∼11 nm. We attribute this difference to intrinsic asymmetries in the three-helix bundle design (**Figure S1 A,C**), as it is not possible to have a perfectly symmetric arrangement of strand crossovers in the constant region for such a simple nanostructure. The three arms were splayed outwards, due to the electrostatic repulsion between the double-stranded DNA arms and the flexible hinges between the arms and the Fc-mimetic bundle. The spatial separation of the nano-synbody trivalent binding domains was within 2 nm of the geometry with three RBD domains in the up configuration, highlighting the agreement with the structure obtained from the computational design pipeline. The three LCB1 proteins are free to rotate due to their flexible attachment to the DNA arms via aliphatic linkers. The 15-atom linkers between the DNA 5’-terminus and the attachment site to the protein surface was parameterized using an all-atomistic model, enabling a realistic simulation of both the distance between the LCB1 proteins and the dsDNA arms, and the relative flexibility and fluctuations thereof.

For the bound structure, we were particularly interested as to whether the nano-synbody could bridge multiple conformations of the spike trimer, namely two RBD domains up and one domain down (**Figure 2D**) and all three domains up (**Figure 2E**). For the former case, the inter-RBD distances vary from ∼3.5 nm for the nearby up and down domains, and 7-8 nm between the distant up and down domain and two up domains (**Figure 2F**), whereas for the more symmetric three-up arrangement, all three distances are approximately equal at 6.5-7 nm (**Figure 2G**). Analysis of the root mean square fluctuation (RMSF), show that the LCB1 proteins at the tips of the nano-synbody arms are easily able to bridge both of these scenarios, with the majority of the structure remaining rigid and only the tips of the arms fluctuating more than 1.8 nm. The nick sites where the three arms meet the three-helix bundle core impart flexibility to the arms, enabling them to adapt to the distances on the spike protein without incurring any strain in the structure relative to the unbound nano-synbody, as seen from the respective RMSF values. This arrangement is similar to IgG molecules, where the hinge regions between the Fc and Fab domains allow the Fv domains to span a range of distances.^37^ As such, the structure balances rigidity for the Fc-mimetic domain and for each arm duplex, with sufficient flexibility between the Fc- and Fab-mimetic domains to accommodate various conformational states of the spike trimer. Based on these results, we anticipated that the hybrid nano-synbody should bind to the spike trimer without incurring a significant entropic penalty due to confining the structure upon binding. Movies showing the fluctuations of the bound and unbound nano-synbody can be found in the online Supporting Information.

### Synthesis of nano-synbodies with one, two, or three arms

In order to modify LCB1 with a ssDNA handle, we introduced a unique cysteine residue at the N-terminus. The protein was expressed recombinantly in *E. coli*, and conjugated to a 14-nucleotide, amine-modified DNA handle using the hetero-bifunctional linker sulfosuccinimidyl 4-(N-maleimidomethyl)cyclohexane-1-carboxylate (SMCC), **Figure 3A**, **S6**, **S7A**. Following exposure to a 1:1 ratio of purified SMCC-DNA, we observed successful conjugation of the LCB1 mini binder to the oligonucleotide, resulting in LCB1-DNA with approximately 90% yield (**Figure S7B**). The modified protein-DNA conjugate was purified using His-trap affinity chromatography to remove excess DNA, followed by anion exchange chromatography to remove excess LCB1 (**Figure S7C**). Denaturing polyacrylamide gel electrophoresis (SDS-PAGE) analysis confirmed the purity of the DNA-LCB1 conjugate, which showed a single band with slower mobility compared with the unmodified LCB1 protein (**Figure 3A**). For the details of protein production, purification, and modification with DNA see the Supporting Information, **Section S1**.

**Figure 3:**
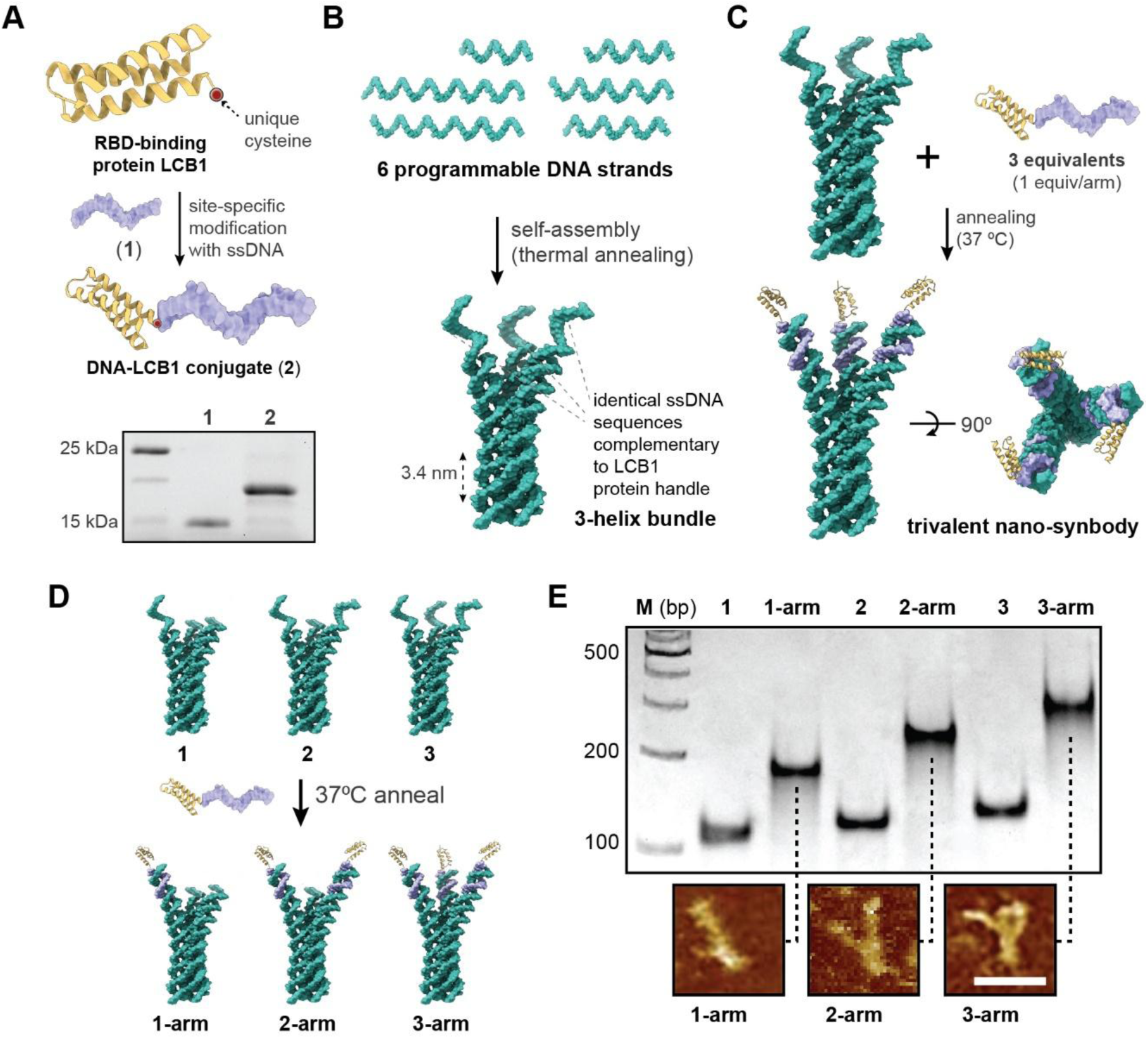
Synthesis and characterization of nano-synbodies with varying numbers of arms. **A)** The LCB1 mini-binder protein was site-specifically modified with DNA using a unique, mutagenically introduced cysteine residue. The gel (SDS-PAGE) shows the shift in retention between the unmodified DNA and the purified DNA-LCB1 conjugate. **B)** The three-helix bundle (3HB) DNA scaffold was annealed from six unique DNA oligonucleotides. **C)** Incubating the 3HB with the DNA-LCB1 conjugate results in a trivalent nano-synbody. **D)** Three-helix bundles with one (**1**), two (**2**), or three (**3**) ssDNA arms were annealed with 1-3 equivalents, respectively, of DNA-LCB1 to yield **1-arm**, **2-arm**, and **3-arm** nano-synbodies. **E)** Native PAGE analysis of the structures from (**D**), showing a shift with increasing arms (lanes **1**, **2**, and **3**) and between samples with and without protein (e.g. **1** vs. **1-arm**, etc.). AFM images show the nano-synbodies with the expected number of arms; to enhance contrast, the samples were incubated with monomeric RBD prior to imaging. Scale bar: 20 nm.

Next, we prepared the 3HB nano-scaffold with three ssDNA arms from six constituent strands via a thermal annealing protocol (**Figure 3B**). Combining the DNA-LCB1 conjugate with the 3HB nanostructure in a 3:1 ratio (so 1:1 equivalent per binding arm), and incubating at 37 °C for 30 min, to allow the DNA to hybridize yields the protein-DNA nano-synbody, which we term “3-arm” (**Figure 3C, S1**). One key advantage of DNA as a nanoscaffold is that, unlike native IgG molecules, the number of Fab-mimetic arms on the nano-synbody can be readily tuned, simply by truncating one or two of the constituent strands. We thus designed two additional nano-synbodies with only one or two complementary handles to DNA-LCB1 (which we term “1-arm” and “2-arm”, respectively, **Figure 3D, S2, S3**) in order to directly compare the effect of mono- and bi-valency of the structure with the tri-valent 3-arm nano-synbody. The 3HB scaffolds with one, two, and three arms could all be assembled readily from the constituent strands, and showed increased retention by native PAGE (**Figure 3E**). Adding DNA-LCB1 to each of these DNA scaffolds resulted in the 1-arm, 2-arm, and 3-arm structures. Native PAGE confirmed that the addition of DNA-LCB1 shifted each nano-scaffold to a higher retention, and the addition of each additional LCB1 protein shifted the final nano-synbody relative to the samples with fewer arms. All samples assembled in high yield (>95%), without aggregation, and did not require additional purification before use. Indeed, the simplicity of our design—which uses the minimum number of strands necessary to recapitulate the desired geometry and distances—will aid in scalable production of the desired nano-synbody. The 1-, 2- and 3-arm nano-synbodies were imaged by AFM after addition of monomeric RBD protein, in order to bind the tips of the arms and enhance contrast. The images clearly showed rod-like structures with 1-3 arms, and dimensions approximately 20 x 4 nm, which match the expected dimensions of the structure (**Figure 3D, S8**).

### Probing multivalent nano-synbody binding to the spike protein trimer

To further characterize the binding interactions between the nano-synbody constructs and the SARS-CoV-2 spike protein, we performed surface plasmon resonance (SPR) measurements using a home-built SPR system based on the Kretschmann configuration (**Figure S10**).^38^ Carboxyl-functionalized gold sensor surfaces were activated using N-hydroxysuccinimide (NHS) and 1-ethyl-3-(3-dimethylaminopropyl)carbodiimide (EDC) chemistry, enabling covalent immobilization of the spike protein via amide bond formation. We first evaluated the binding affinity of the purified LCB1 protein toward the wild-type (WT) and Omicron BA.1 spike proteins, serving as high- and low-affinity targets, respectively (**Figure 4A** and **4B**). LCB1 bound to the WT spike protein with an equilibrium dissociation constant (K_D_) of 1.22 nM, which was on par with the value in the original report of this protein.^27^ In contrast, no detectable SPR response was observed for the Omicron spike protein, indicating a dramatic loss of binding. This result is consistent with the fact that LCB1 was originally designed to recognize the WT receptor-binding domain (RBD), and that multiple mutations present in the Omicron variant substantially alter the epitope recognized by the mini-binder. Next, we examined whether conjugation of LCB1 to DNA affected its binding capability. The DNA–LCB1 conjugate displayed a binding affinity to the WT spike protein of 1.64 nM, comparable to that of free LCB1, while still showing no measurable interaction with the Omicron spike protein. These results indicate that DNA functionalization does not compromise the structural integrity or binding functionality of LCB1. Similarly, the monovalent (1-arm) nano-synbody, consisting of a single DNA–LCB assembled onto a three-helix bundle (3HB) scaffold, retained a binding affinity to the WT spike protein of (1.18 nM) and no detectable binding to the Omicron spike, results that are comparable to that of free LCB and DNA–LCB. It was somewhat surprising that a large 3-helix bundle nanostructure appended to LCB1 had no deleterious effect on its binding to the WT protein, but we attribute this to the long and flexible linker between the protein and the DNA. As evidenced in our simulations (see movies in the Supplementary Information), the linker allows the LCB1 significant conformational freedom; all-atom MD yields a linker length of ∼1.8 ± 0.8 nm (modeled as a skewed Gaussian).

**Figure 4:**
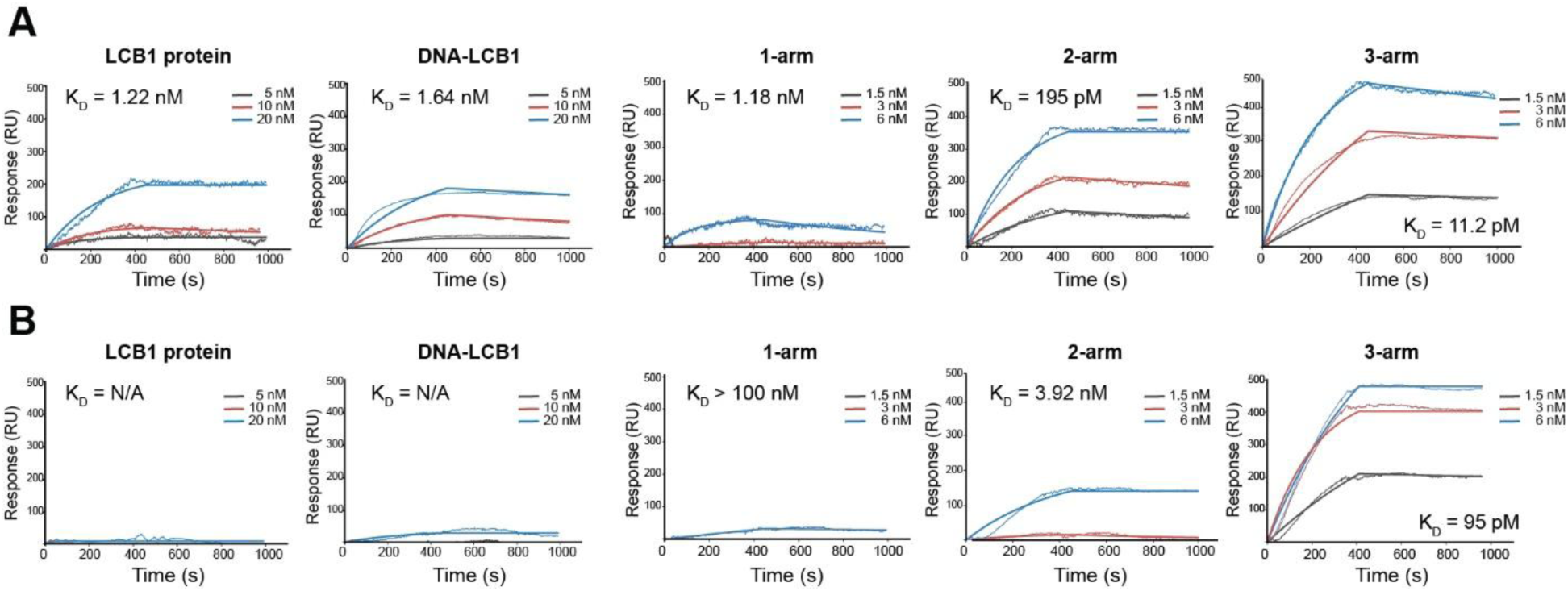
Surface plasmon resonance characterization of nano-synbody affinity. SPR sensorgrams of indicated species, for the wild-type protein (**A**), and the Omicron BA.1 variant (**B**). LCB1 is the mini-binder protein alone; DNA-LCB-1 is the ssDNA-protein conjugate; 1-, 2-, and 3-arm correspond to the samples from Figure 3D. The equilibrium dissociation constant for each sample (K_D_) is indicated on the graph.

We then systematically compared the binding properties of mono-, bi-, and trivalent nano-synbody constructs against both WT and Omicron spike proteins (**Figure 4A,B**). For the WT spike protein, a clear valency-dependent enhancement in binding affinity was observed. Compared to the 1-arm nano-synbody, the 2-arm nano-synbody showed an improved affinity, with a K_D_ of 195 pM, corresponding to approximately a 6-fold enhancement relative to the monovalent construct. The 3-arm nano-synbody achieved an affinity with a K_D_ of 11.2 pM, representing over a 100-fold improvement compared to the monovalent configuration, and a 17-fold improvement over the 2-arm structure. This enhancement is indicative of strong avidity effects arising from multivalent engagement of the spike protein. Notably, multivalency also enabled effective binding to the Omicron BA.1 spike protein, despite the negligible interaction observed for free LCB1 and the monovalent nano-synbody (K_D_ > 100 nM for the 1-arm structure). The bivalent (2-arm) nano-synbody exhibited measurable binding to the Omicron spike protein, with a K_D_ of 3.92 nM, while the trivalent (3-arm) nano-synbody demonstrated a strikingly high binding affinity of 95 pM, a roughly 40-fold enhancement over the 2-arm structure, and at least a 1000-fold enhancement over the monovalent structure. These results suggest that multivalent presentation of LCB1 can compensate for weakened individual binding interactions caused by viral mutations. **Table S1** summarizes the SPR data, including association and dissociation rates.

These SPR studies clearly show that the relative increase in affinity for the nano-synbody is markedly greater for the Omicron variant than the WT protein. Our results are in line with a number of other studies that multimerize spike-targeting protein binders,^30,39^ whereby a greater enhancement in relative binding affinity was seen for mutated spike proteins that were poorly targeted by monovalent binders. Other systematic examinations of multivalency have also found that weaker monovalent binders benefit from a bigger boost due to avidity effects, and yield a greater selectivity of binding.^40–43^ However, we note that some studies have found the inverse, where a higher monovalent binding affinity yields a greater multivalent enhancement than a weaker binder.^44^ This discrepancy may be due to the specific scaffolds, geometries, reconfiguration penalties, and assays used in the respective systems, which makes a direct comparison of systems difficult. Nonetheless, the ability to systematically tune the number of ligands, or the geometry and rigidity of the nano-scaffold (both of which are possible with the nano-synbody) will allow for fruitful investigations of the various factors driving avidity in future studies.

### Nano-synbody neutralization of pseudovirus infection

We next investigated whether the multivalency enhancement observed in our SPR studied could translate to improved SARS-CoV-2 neutralization, using a pseudovirus assay (**Figure 5A**). We engineered pseudovirus particles (according to a reported protocol)^45^ bearing spike proteins for the WT virus and three Omicron sub-variants (BA.1, BA.2, and BA.4/5). The pseudovirus contained either a GFP or a luciferase gene, allowing successful infection to be monitored using either fluorescence or bioluminescence, respectively. Pseudovirus infection was probed using a human embryonic kidney (HEK) 293T cell line transfected with hACE2 receptors; for cell line and pseudovirus characterization see **Figure S11,12**. We first probed virus neutralization using the GFP gene-bearing pseudovirus. Cells were exposed to the pseudovirus (1 × 10^6^ TU/mL functional titer; TU = transduction units) and 10 nM of the 3-arm structure. As seen in **Figure 5B,C**, in the absence of the nano-synbody, cells exposed to pseudovirus bearing the WT or Omicron BA.1 spike trimers showed cells with extensive green fluorescence, indicating successful infection. Cells lacking the hACE2 receptor did not show any fluorescence (see **Figure S11D**). By contrast, addition of the 3-arm nano-synbody resulted in virtually no GFP fluorescence for either the WT or Omicron BA.1 spike proteins, confirming that at this concentration it could almost completely block pseudovirus infectivity.

**Figure 5:**
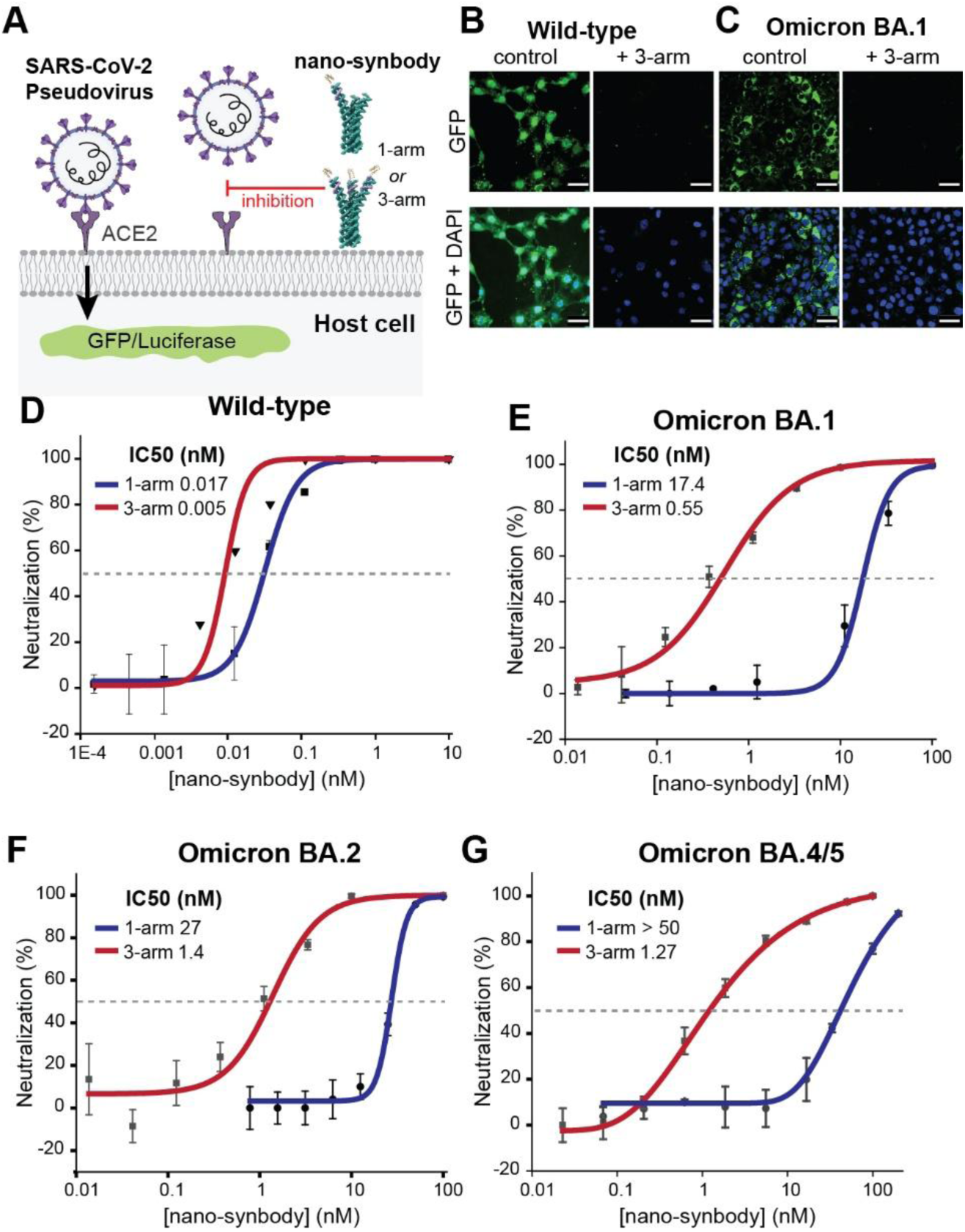
Pseudovirus neutralization assay. **A)** Schematic representation of the neutralization assay for the SARS-CoV-2 pseudovirus by the nano-synbody. **B,C)** Confocal fluorescence images depicting pseudovirus infection (with GFP cargo) with different spike variants in the presence or absence of 3-arm nano-synbody (10 nM). Scale bar: 50 μm. **D-G)** Demonstrated neutralization of the indicated SARS-CoV-2 pseudovirus variants by the nano-synbodies (luciferase cargo). The spike pseudoviruses were subjected to co-incubation with serial dilutions of both 1-arm and 3-arm nano-synbodies for 30 minutes. Subsequently, the mixture was introduced to 293T-ACE2 cells. Following a 36-hour period, the IC_50_ values for 1-arm or 3-arm nano-synbodies were determined by assessing luciferase expression levels in the 293T-ACE2 cells. Data are representative of three independent experiments. Mean ± SD was shown.

For a more quantitative and sensitive measurement of pseudovirus neutralization, we turned to a luciferase-based assay and quantified IC_50_ values for the nano-synbodies. Given our previous results, we decided to only compare the 1-arm and 3-arm nano-synbodies in this assay, to determine the maximum enhancement that multivalency could provide. Similar to the SPR results in **Figure 4**, the nano-synbodies showed the greatest neutralization for the pseudovirus bearing the WT spike, with IC_50_ values of 5 and 17 pM for the 3-arm and 1-arm structures, respectively (**Figure 5D**). This roughly 3-fold change due to the multivalency was also the smallest difference for all the variants tested. However, several IC_50_ values approached the assay’s operational lower limit of quantification (LLOQ; ∼5 pM, defined by the lowest concentration tested)^46^ so we cannot determine an affinity enhancement beyond this point. By contrast, the IC_50_ values for the 3-arm vs. 1-arm nanostructures were 548 pM vs. 17.4 nM for the Omicron BA.1 spike (∼30-fold enhancement, **Figure 5E**); and 1.4 vs. 27 nM for the Omicron BA.2 spike (∼20-fold enhancement, **Figure 5F**). For the Omicron BA.4 spike, only the 3-arm nano-synbody gave a measurable IC_50_ value (1.3 nM); the 1-arm structure never reached full inhibition (i.e., IC_50_ > 50 nM; so the 3-arm structure demonstrated >40-fold enhancement), **Figure 5G**. Taken together, these results parallel our findings by SPR, enabling multivalent inhibition even of variants for which the monovalent ligand is ineffective.

### Structural characterization of the 3-arm nano-synbody to the Omicron spike trimer

One key advantage of our approach is that the rigid Fc-mimetic 3HB could serve as a rigid handle to integrate the protein with other surfaces or nanostructures, e.g. a DNA origami scaffold, in order to immobilize and/or manipulate the bound target. We thus sought to characterize the 3D structure of the nano-synbody bound to the spike trimer by negative-stain (ns) TEM, and specifically to show that all three arms were attached as designed. The 3-arm structure showed almost complete binding to the Omicron spike trimers at a 1:1 stoichiometric ratio, as evidenced by agarose gel electrophoresis (**Figure S9**). We next analyzed the Omicron spike trimer (B.1.1.529) both before and after binding to the 3-arm nano-synbody, via nsTEM. We determined the structure to ∼20 Å resolution by 3D reconstruction via reference-free 2D classification using the Relion3.1 software; see **Section S1** for the details of sample preparation, TEM image acquisition, and 3D reconstruction. The spike protein alone showed a conformation with one RBD in the “up” position, which aligns with previous high-resolution cryo-electron microscopy characterizations (**Figure 6A,B**).^47,48^ The single RBD in the up conformation was not well resolved due to structural flexibility compared to the rest of spike trimer. Despite the relatively low resolution of our structure, we were able to distinguish the three individual spike proteins comprising the trimer, including the two RBD domains in the “down” position, characteristic of the virus prior to binding the ACE2 receptor.

**Figure 6:**
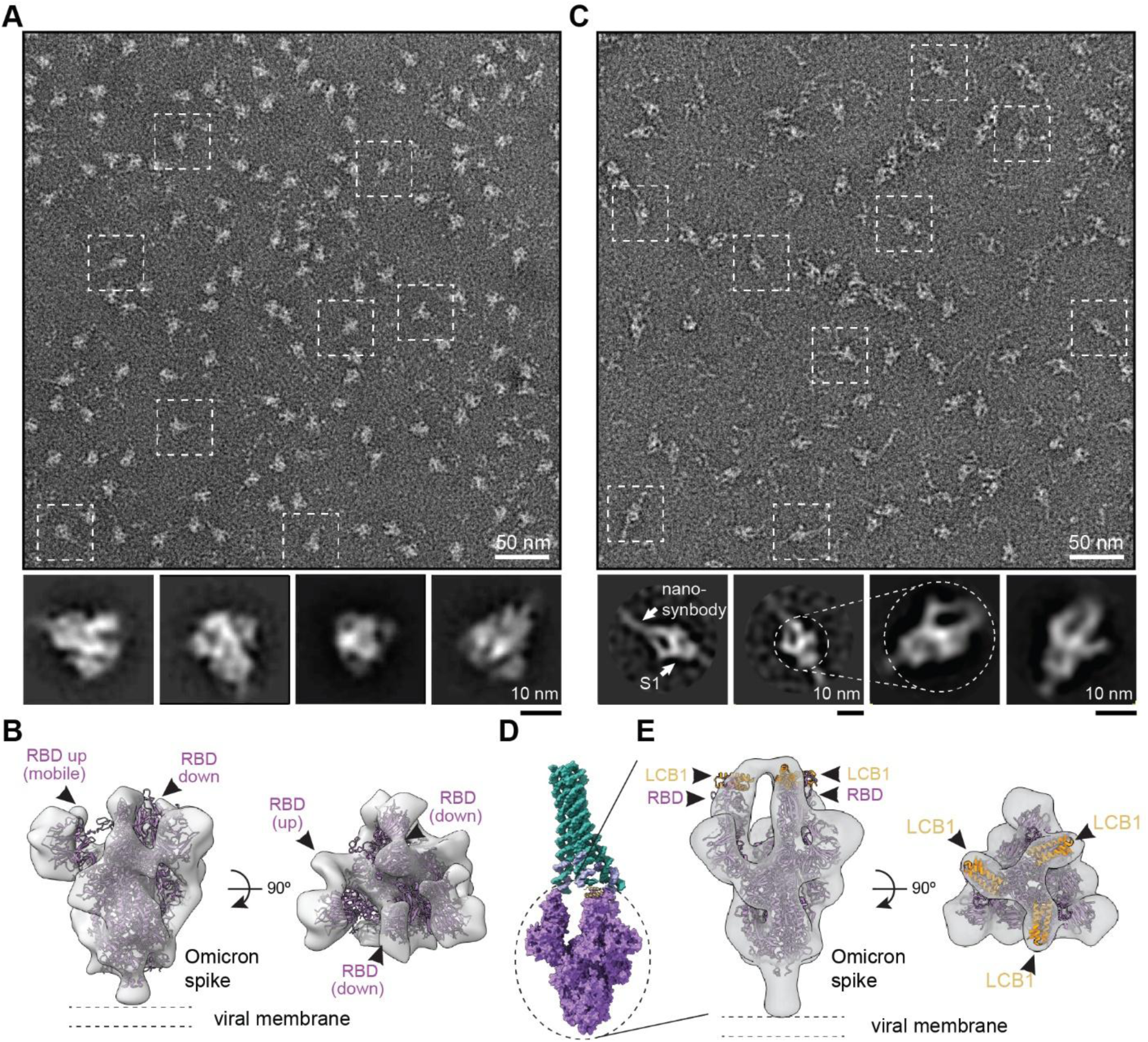
Negative-stain (ns) TEM structural characterization of nano-synbody binding to the Omicron spike trimer. **A,C)** Representative nsTEM images of the Omicron B.1.1.529 spike trimer before (**A**) and after (**C**) exposure to the 3-arm nano-synbody. The smaller images show representative 2D class averages for the samples. In (**C**), the 3HB core of the nano-synbody can be distinguished, along with multiple arms binding to the S1 proteins of the spike trimer. **B)** A negative-stain 3D reconstructed EM map (*gray*) of spike trimer in two orthogonal orientations. A cryo-EM map (*violet*) of the spike trimer (PDB: 7THK) was docked into the EM map for visualization. **D)** Representative illustration of nano-synbody (*green*) bound with spike trimer (*purple*) with three RBDs in the up conformation. **E)** A focused negative-stain 3D reconstructed EM map of the nano-synbody complex with the spike trimer in two orthogonal conformations. The EM map shows the spike trimer with three RBD in the up conformation and bound to LCB1.

By contrast, the 2D class averages of the Omicron spike trimer exposed to the 3-arm nano-synbody showed additional density corresponding to the cylindrical nano-synbody (∼10 nm in length), connected with three arms to the S1 domains of the spike trimer (**Figure 6C,D**). Although the spike protein could be reconstructed (similar to **Figure 6B**), the fluctuations of the DNA nanostructure—resulting from the flexible linkages between the 3HB core of the structure and the three duplex arms, as well as the flexible chemical linkers between the ends of the arms and the LCB1 protein—precluded reconstruction of the full nano-synbody bound to the spike protein. However, both the 3HB core and the arms are visible in the 2D class averages, indicating that the structure was intact and bound as designed. We applied a smaller circular mask that included the S1 domain of the spike trimer and the region where three LCB1 proteins were designed to bind with RBD, while excluding most of the nano-synbody structure (as shown in the 2D class images in **Figure 6D,E**). The 3D reconstruction map was generated using these 2D classes, which allowed the atomic model of the spike protein (PDB 7UHC)^30^ to be docked successfully into the map using rigid body fitting in ChimeraX. In this 3D reconstructed image, we were able to observe density corresponding to the three RBD domains in the up position, bound to three copies of the LCB1 protein (**Figure 6E**), matching the reported structure of the monomeric LCB1 bound to RBD by cryo-EM.^30^ Furthermore, all three RBDs were resolved in the up position, with distances of ∼7 nm between them (corresponding closely to the simulated model, **Figure 2E,G**). In particular, we clearly resolved three copies of LCB1 bound to the trimer, strongly suggest that all three arms of the nano-synbody are competent to engage with the Omicron spike trimer as designed. Future studies at higher resolution using cryo-EM should be able to further explore the structure of the protein-bound DNA scaffold itself.

### Reversible protein binding through dynamic valency switching

Beyond affinity enhancement, a key advantage of the nano-synbody platform is its ability to enable precise and reversible control over binding valency. While multivalent effects are widely exploited in many nanostructure-based systems, achieving exact and dynamically tunable valency in these platforms remains challenging. In contrast, the DNA-based architecture of our nano-synbody allows programmable regulation of binder valency with single-arm resolution. To demonstrate reversible valency control, we employed toehold-mediated strand displacement^49^ to selectively remove DNA-LCB1 binding arms from the trivalent (3-arm) nano-synbody. Specifically, the 3’ ends of the DNA-LCB1 binding handles were extended with three different 10-nucleotide toehold domains (labeled a, b, and c (**Figure 7A, S4**)). These toeholds serve as initiation sites for complementary invader strands, enabling controlled displacement of DNA-LCB1 conjugates and conversion of the nano-synbody from trivalent to the bivalent, monovalent, or “zero-valent” (i.e. the 3HB with no proteins attached) states. We highlight that, because the displacement strand replaces the DNA-LCB1 conjugates, leaving behind a DNA duplex devoid of any protein at its end, these displaced states are not exactly identical to the 2-arm, 1-arm, or bare 3HB structure from **Figure 3D**, which lack the specified number of arms altogether.

**Figure 7:**
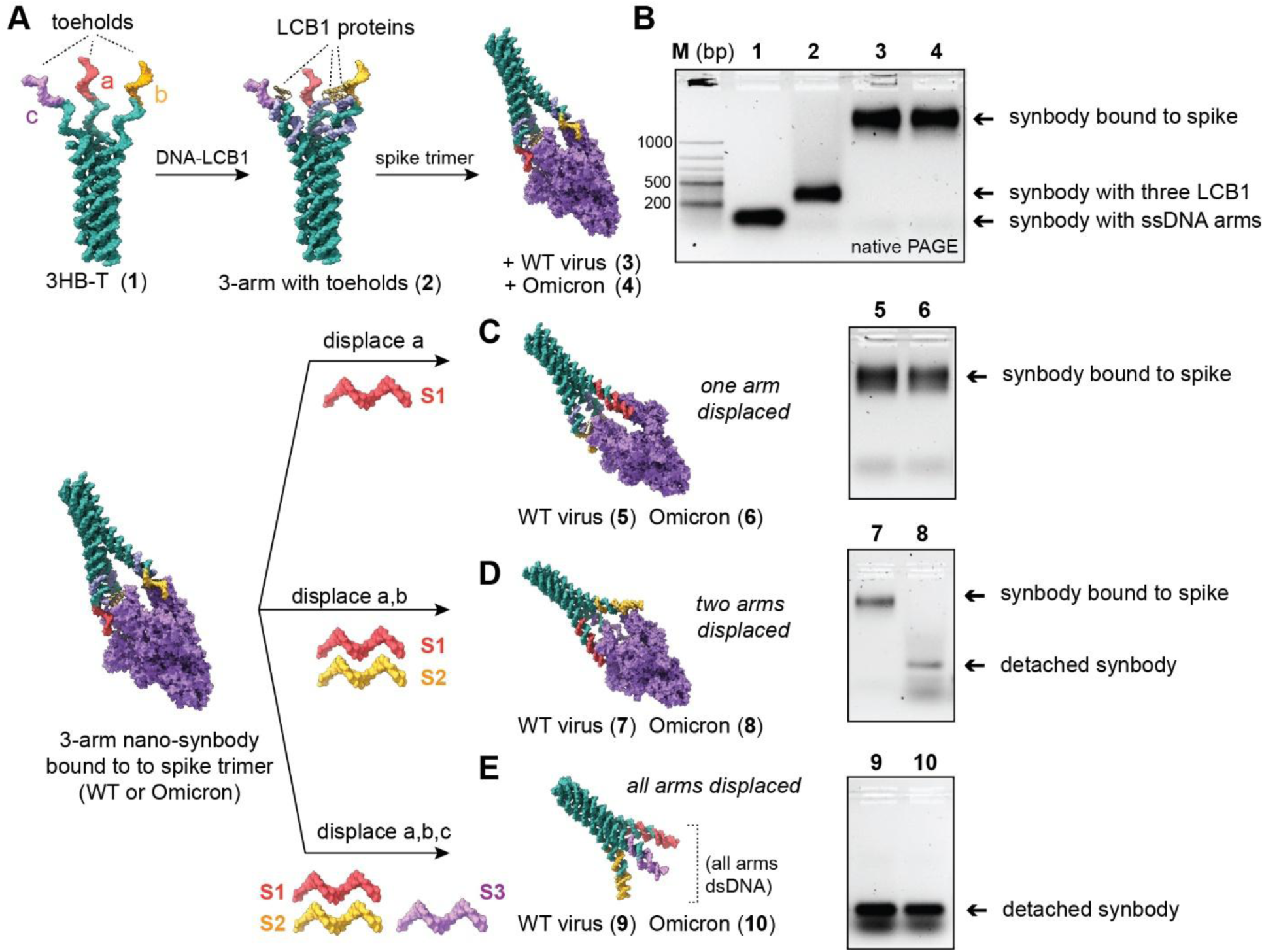
Programmable reversal of multivalent binding. **A)** Schematic of nano-synbody with three different arms, each bearing a unique toehold sequence (a, b, and c). Addition of DNA-LCB1 generates a 3-arm structure, with three ssDNA toeholds (purple, red, yellow). **B)** Native PAGE of this 3-arm nano-synbody bound to the spike trimer. The lanes indicate the species from (**A**). Complete binding of both WT and Omicron spike trimers can be seen by the 3-arm nano-synbody (**2**). **C-D)** Toehold mediated displacement of one (**C**), two (**D**), or three (**E**) arms of the synbody. Native PAGE gels indicate the numbered species after the displacement, for both the WT and Omicron variants.

The 3-arm nano-synbody bearing all three toehold domains was first assembled and incubated with either the WT or the Omicron spike trimers. A clear and complete shift to a higher molecular weight species corresponding to the bound complex can be seen by PAGE (**Figure 7B**), due to the picomolar affinity of the 3-arm structure for both protein variants. Next, we incubated the bound nano-synbody/spike complex with one, two, or three specific invader strands (S1, S2, and S3; which replace toeholds a, b, and c, respectively) at 37 °C for 10 min. Introduction of a single invader strand resulted in displacement of one DNA-LCB1 arm, producing a bivalent nano-synbody (**Figure 7C**). Addition of two invader strands in combination (S1+S2, S1+S3, or S2+S3) displaced two DNA-LCB arms, yielding a monovalent construct (**Figure 7D**). Complete removal of all three DNA-LCB1 units was achieved by adding all three invader strands simultaneously (S1+S2+S3), regenerating the three-helix bundle (3HB) scaffold, albeit with three dsDNA arms (**Figure 7E**). Optimization of reaction stoichiometry showed that a 1:5 molar ratio of 3-arm nano-synbody to the combined invader strands, for 10 min at 37 °C, was sufficient to achieve complete displacement (**Figure S14**). To assess how reversible valency control modulates spike protein binding, we performed native gel-based binding assays.

The 3-arm nano-synbody (200 nM) was incubated with either WT or Omicron spike protein, followed by treatment with different combinations of invader strands. Following displacement of a single DNA-LCB arm, the resulting bivalent nano-synbody retained strong binding to both WT and Omicron spike proteins (**Figure 7C**). This result is consistent with the SPR measurements, which show picomolar and low-nanomolar affinities, respectively, well below the concentration of the synbody added. In contrast, conversion to a monovalent nano-synbody led to selective binding: the WT spike protein remained tightly associated, whereas the Omicron spike protein was no longer captured, as indicated by the disappearance of high-molecular-weight bands and the appearance of a band corresponding to the free nano-synbody structure (**Figure 7D**). Once again, this is consistent with the SPR results, where the monovalent nano-synbody still showed low-nanomolar affinity for the WT virus, but no measurable affinity for the Omicron variant. Complete displacement of all three DNA-LCB arms resulted in release of both spike variants, leaving only free nano-synbody scaffolds and displaced DNA strands (**Figure 7E**). Together, these results demonstrate that toehold-mediated strand displacement enables precise and reversible regulation of nano-synbody valency, thereby providing dynamic control over protein binding.

Such a strategy allows not only modulation of binding strength but also selective engagement and release of target proteins in response to programmable molecular cues, highlighting the versatility of DNA-assembled nano-synbodies as adaptive and reconfigurable protein-binding platforms. Moreover, although all three toehold-bearing strands in **Figure 7A** are different, in order to enable for systematic removal of 1-3 LCB1 proteins, making all three toeholds identical would allow for complete release of both WT and Omicron variant trimers with three equivalents of a single displacement strand. Although the LCB1 proteins, after displacement, still presumably remain bound to the WT spike (due to their ∼1-nM K_D_), the extremely poor affinity for the Omicron variant means that the now-monovalent protein should dissociate easily. As such, we believe that the nano-synbody can serve to not only turn a bad binder into a good one, through multivalency, but effectively completely abolish target inhibition by reverting to the low-affinity ligand on-demand. This is, to our knowledge, one of only a few examples^50–52^ that use DNA-mediated displacement to control polypeptide binding to molecular targets, and the first to systematically reverse valency between multiple states, for multiple targets of varying affinity.

## DISCUSSION

We have demonstrated that an IgG-mimetic DNA nano-scaffold can be designed to match the geometry and nanoscale spacing of the SARS-CoV-2 spike trimer, displaying three copies of an RBD-binding mini-protein. We used a computational pipeline incorporating the coarse-grained, oxDNA-ANM model to engineer the hybrid nanostructure, and to minimize the number of strands necessary. This nano-synbody was able to bind the wild-type and Omicron variants of the spike protein, and neutralize pseudovirus infectivity by blocking the RBD interaction with the ACE2 receptor. The modularity and programmability of the DNA scaffold enabled direct comparison of nano-synbodies with one, two, and three arms; the three-arm structures were shown to have the highest binding affinity (and concomitant neutralization potential), with multivalency playing an increasingly important role for variants with mutations that weakened the monovalent LCB1-RBD interaction. For several Omicron variants, the 3-arm nano-synbody showed both efficient binding and neutralization, highlighting the power of multivalency to “rescue” a compromised monovalent binder. Moreover, we could systematically and programmably reverse the binding by displacing one, two, or all three arms of the nanostructure after binding to the spike protein, allowing for binding and unbinding through stepwise, “bidirectional” multivalency. Our system is simple, consisting of only six DNA strands and a single protein-DNA conjugate, and can be carried out in high yield and efficiency while enforcing a nanoscale spacing between the three protein ligands that can match multiple spike trimer conformations. As such, our nano-synbody is more complex than a DNA duplex linker, but much simpler than a DNA origami nanostructure, which can consist of ∼200 oligonucleotides. The relative simplicity of the nano-synbody design will allow for modified nucleotides to be used in the future, if necessary, to confer stability and/or reduce immunogenicity for *in vivo* applications.

The concept of trimerization to match the spike protein has been employed extensively by nanotechnological approaches to SARS-CoV-2 neutralization. Indeed, both LCB1^30^ and a number of other protein-based viral binders^53–58^ have been oligomerized using either linear or branched linkers in order to increase the binding affinity. These systems also showed enhanced binding due to multivalency, as well as resistance to mutational escape for multiple variants. Compared with these all-protein based approaches, a protein-DNA nano-synbody has several advantages for certain applications. First, DNA nano-scaffolds enable matching the size, shape, and valency of a target to a higher degree than a protein-based scaffold. It is relatively straightforward to tune the size of the rigid DNA bundle (e.g. expand from three to six,^59^ twelve,^60^ 24 helices,^61^ or even larger bundles), or include multiple *different* binding agents (since each arm can be extended with a unique DNA handle) in a highly precise and stoichiometrically-defined fashion. As such, a DNA nanostructure could in principle bind to patches away from a key interface, which are less likely to mutate, and block protein-protein interactions by dint of the bulky nanostructure. Indeed, nanobodies that bind to the N-terminal domain (NTD) of the spike protein have been reported,^62^ and we are currently exploring creating a heterobivalent nano-synbody that positions one of these nanobodies along with LCB1 to enhance binding to a single spike monomer. Our design pipeline can also readily generate nano-synbodies that span distances of ∼10-50 nm, lengths scales that are challenging for protein-based approaches, but for which origami-based systems might be unnecessarily large and DNA-uneconomical.

Furthermore, our nano-synbody approach can, in principle, easily integrate multiple types of binding groups, combining proteins with aptamers, peptides, or other molecular species like carbohydrates.^63^ The rigid, Fc-like domain can serve additional functions, such as attaching: imaging agents; stoichiometrically-precise numbers of cytotoxic agents for antibody-drug conjugate (ADC) mimics; polymers to prolong circulation in vivo or prevent protein fouling; other effector proteins; or oligonucleotide handles for detection methods like rolling chain amplification.^64^ hairpin chain reaction,^65^ or polymerase exchange reaction.^66^ Although the 3HB core in this work was designed to require as few strands as possible, the length could be extended in order to tune the number and/or spacing of attached molecules, like biological Fc domains to interface with the immune system. The modularity of our approach should also allow for new binders—such as “updated” versions of LCB1,^30^ or nanobodies^67,68^ that can more effectively bind to the Omicron spike—to be swapped in, without requiring redesign of the core structure as long as the target spacing remains unchanged. Moreover, the Fc-mimetic domain can also serve as a structured handle to further assemble the nano-synbodies into hierarchical structures using branched linkers, e.g. to yield IgM-mimetic structures but with valencies beyond the classic pentamer. Branched DNA junctions can attach anywhere from three^69^ to twelve^70^ nano-synbodies by extending an addressable handle from the rigid domain. If the hierarchically-assembled nano-synbodies bind to two or three *different* targets, it should be possible to create synthetic bi- and tri-specific antibodies^71–73^ with a high degree of tunability and modularity, in both the valence and the spacing between the cells targeted.

Finally, our system is unique compared with other multivalent, spike-targeting nanosystems in that we demonstrate valency reduction through selective removal of one, two, or three LCB1 ligands. Such effectively “reversible IgG” mimics could be used as blocking antibodies that can be deactivated on demand, turning “on” the activity of their target. Because DNA toeholds are sequence specific, multiple blocking nanostructures can be used in parallel, and each one disabled when desired, using specific displacement strands. These systems can also serve as fluorescently-tagged antibodies that can be erased when desired by adding displacement strands, to enable multiple rounds of staining and imaging. By demonstrating that the affinity could be tuned through both the monovalent binder strength, and the number of arms, our approach could be used to enhance specificity by positioning multiple weak binders with spatially matched distances, as demonstrated using a wide range of other ligands.^40–42^

Although in this work we focused on SARS-CoV-2 neutralization as a way to measure the multivalency of binding, our nano-synbody can be used for applications beyond biomolecular targeting, imaging, or detection. For instance, the integration of a multivalent binding interface with a rigid, nanostructured domain would allow the nano-synbody to act like a nanoscale “gripper arm,” that can bind a target with high affinity and manipulate it at the single-molecule level. The rigid domain could be used to interface with DNA nano-mechanical devices^74^and exert a force or to position them in two- and three-dimensional space using DNA walkers^75^, crank-slider devices,^76^ or 2D “printers.”^75^ In such applications, the ability to displace the protein would allow for a single-molecular manipulator that can pick up a target, move it to a desired location, and release it; the gripper arm could then be recharged by adding fresh protein-DNA conjugates to repeat the process in a catalytic, or potentially fueled/dissipative fashion.

## Acknowledgements

Research reported in this publication was supported by The National Institute of General Medical Sciences of the National Institutes of Health under grant number DP2GM132931 to N.S., grant number 1R01GM145916-01A1 to N.S. and P.S., grant number R01GM140193 to S.W, and grant number R01GM155563 to H.Y. The content is solely the responsibility of the authors and does not necessarily represent the official views of the National Institutes of Health. P.S. acknowledges support from National Science Foundation grant DMR 2239518. N.S. gratefully acknowledges Steve Huffman for a generous gift supporting this research. The authors are grateful to Prof. Fang Li (U. Minnesota) for providing the spike protein variants used.

## Notes

### Competing Interest Statement

The authors have declared no competing interest.

### Summary of Updates

This version of the manuscript has been significantly revised to include new data, as well as new experimental directions.

