## Supplementary Information for "IgG-inspired, multivalent protein-DNA nanostructures for high-affinity, tunable, and reversible binding to biomolecular targets"

**Table of contents** **Pages**


| | |
|---|---|
| Section 1. Materials and Methods | 3 |
| Section 2. Simulations of nano-synbody with and without spike protein | 7 |
| Section 3. Design and DNA sequences used for nano-synbody synthesis | 10 |
| Section 4. Additional characterization data | 15 |
| Section 5. References | 25 |

### S1. Materials and Methods

#### Reagents

All DNA strands were purchased from Integrated DNA Technologies (IDT). Ethyl alcohol (pure, 200 proof), hydrochloric acid (HCl, 37%), 11-mercaptoundecanoic acid, and ethanolamine hydrochloride (ETA·HCl) were purchased from Sigma-Aldrich. N-hydroxysuccinimide (NHS) and 1-ethyl-3-(3-dimethylaminopropyl) carbodiimide hydrochloride (EDC) were purchased from ThermoFisher. The 22 x 22 mm glass coverslip was purchased from VWR. Phosphate buffered saline (PBS) was purchased from Corning. Deionized (DI) water with resistivity of 18.2 MΩ·cm was used in all experiments.

#### LCB1 expression and purification

The LCB1 protein-encoding genes were synthesized and subcloned into pET-28a-TEV plasmid vectors, featuring a modified N-terminal 6×His-tag and a subsequent TEV cleavage site (GenScript). The resultant plasmids were transformed into Lemo21(DE3) *E. coli* strain (New England Biolabs) and cultured in Luria-Bertani medium. Cells were grown in a 2 L baffled flask, supplemented with 50 µg/mL kanamycin, at 37 °C. Upon reaching an optical density (O.D. 600) of 0.6 - 0.8, protein expression was induced with 1 mM isopropyl-β-D-1-thiogalactoside (IPTG, Sigma Aldrich). Following induction, the bacterial culture was incubated for an additional 18 h at 18 °C. Cells were harvested by centrifugation at 10,000 g for 30 min at 4 °C, and the resulting pellets lysed in lysis buffer (25 mM Tris, 300 mM NaCl, 1 mg/ml lysozyme, 0.1 mg/ml DNase, cOmplete™, EDTA-free Protease Inhibitor Cocktail). The lysate was clarified by centrifugation at 20,000 g for 30 min at 4 °C, and the supernatant was filtered through a 0.22 µm syringe filter.

The clarified lysate, which contained soluble LCB1, was subjected to immobilized metal affinity chromatography (IMAC) using a 5 mL HisTrap HP column (Cytiva) on an AKTA pure FPLC system (Cytiva). The column was purged with a washing buffer (1×TBS, 500 mM NaCl, 30 mM imidazole, pH 7.4) before elution of protein fractions using a buffer with 500 mM imidazole in TBS. The eluted fractions were subsequently analyzed by 4-20% stain-free SDS-PAGE precast gels (Bio-Rad) under reducing conditions. The proteins were dialyzed overnight against 1×PBS. The concentration of the purified LCB1 protein was determined by measuring the absorbance at 280 nm, with an assumed extinction coefficient of 9970 $M^{-1}cm^{-1}$. Lastly, the protein was aliquoted (50 µM concentration) and stored at -80 °C in 1×PBS buffer supplemented with 2 mM DTT at pH 7.4.

**LCB1_Cys protein sequence** (introduced Cys for bioconjugation to DNA is underlined):

MSHHHHHHHHENLYFQGGSDKEWILQKIYEIMRLLDELGHAEASMRVSDLIYEFMKKGDERLL
EEAERLLEEVERGS<u>C</u>

#### Synthesis of LCB1-oligonucleotide conjugates (DNA-LCB1)

A 5'-Amino Modifier C6 oligonucleotide handle was procured from Integrated DNA Technologies and dissolved in DNase/RNase-free water (Invitrogen) to achieve final concentration of 1 mM. The amine-DNA was first functionalized with the heterobifunctional linker sulfosuccinimidyl 4-(N-maleimidomethyl) cyclohexane-1-carboxylate (SMCC, Thermo Scientific) as follows: the DNA solution was combined with a 20-fold molar excess of Sulfo-SMCC in 1×PBS (100 mM sodium phosphate, 150 mM NaCl, pH 7.2) buffered solution and dry DMSO. The reaction mixture was incubated at RT, under agitation at 1000 rpm, for 2-4 h. The SMCC-functionalized oligonucleotide was subsequently isolated via ethanol precipitation, which involved the addition of a 300% (v/v) volume of ice-cold ethanol followed by a 45-min incubation at -80 °C. After centrifugation at 20,000 g for 20 min at 4 °C, the supernatant was decanted, and the pellet was re-dissolved in 1×PBS.

This precipitation step was performed twice for maximum purification. The pellet was then resuspended in 1×PBS and washed three times using a 3 kDa molecular weight cutoff (MWCO) Amicon Ultra centrifugal filter (Millipore). The functionalization process was monitored by matrix-assisted laser desorption/ionization time-of-flight mass spectrometry (MALDI-TOF MS). For the conjugation of LCB1 to the SMCC-functionalized DNA handle, LCB1 aliquots were treated with 2 mM DTT to ensure reduction of any disulfide bonds, to yield the LCB1 monomer. This reduced mixture was buffer exchanged into a 100 mM sodium phosphate buffer (pH 7.0) using PD10 desalting columns (GE Healthcare). The desalted LCB1 solution was mixed with a three-fold molar excess of maleimide-functionalized DNA, and the reaction was incubated at 300 rpm overnight at 4 °C. The conjugation efficiency was evaluated using non-reducing SDS-PAGE (**Figure S5B**).

**Purification of DNA-LCB1 conjugates**

The purification of the DNA-LCB1 conjugate was accomplished using a two-step strategy employing fast protein liquid chromatography (FPLC, ÄKTA Pure, GE Healthcare). First, an anion-exchange HiTrap Q HP column (1 mL, GE Healthcare) was equilibrated with a 50 mM Tris-HCl buffer (pH 7.5) by running 10 mL of the buffer through the column. The conjugation reaction mixture was then manually loaded onto the equilibrated column. After loading, the column was re-equilibrated with 15 mL of the equilibration buffer, and a linear salt gradient was applied with starting and ending concentrations of 100 and 500 mM NaCl, respectively, at a flow rate of 1 mL/min. The resulting elution fractions were collected and analyzed via SDS-PAGE to ascertain the separation of unconjugated LCB and LCB1-DNA.

For the second purification step, $Ni^{2+}$-affinity chromatography was employed to remove unconjugated DNA. The LCB1-DNA fractions obtained from the previous step were loaded onto a 1 mL HisTrap HP column (Cytiva), washed with the same buffer used for the initial LCB1 purification, and eluted with the corresponding elution buffer. The eluted fractions were pooled, and the buffer was exchanged to 1×PBS via overnight dialysis (SnakeSkin, 7 kDa MWCO, Thermo Scientific). The concentration of the purified DNA-LCB1 was determined by measuring the absorbance at 230 nm and referencing this value to a calibration curve obtained from pure LCB1.

**Folding, purification and characterization of 3-helix bundle and nano-synbody.**

***ssDNA strand purification.*** All ssDNA strands (sequences listed in **Figure S1B**, **S2B**, and **S3B**) purchased from IDT were purified using an in-house prepared 10% denaturing polyacrylamide gel electrophoresis with 1×TBE as the running buffer. The desired bands on the gel were excised and placed in an elution buffer, and continuously shaken for 12-24 h at 4 °C. Subsequently, the strands were passed through a 3 kDa Amicon filter to remove salts and other soluble impurities. This step was repeated 5-6 times, with each repetition involving supplementation of the supernatant with fresh milli-Q water. Finally, all purified strands were stored at -20 °C for further use.

***3-helix bundle formation.*** All 3-helix DNA nanostructures (1-arm, 2-arm, and 3-arm) were prepared by mixing ssDNA strands in equimolar ratio in 100 µL 1×TAE-12.5 mM $MgCl_2$ buffer at a concentration of 1 µM. Thereafter, reaction mixtures were annealed in a PCR thermocycler instrument using following protocol: mixtures were heated at 95 °C for 5 min, followed by a gradient of 88-76 °C at a rate of 1 °C/min, followed by a gradient of 76-24 °C at a rate of 0.5 °C/min and then quickly cooled to 4 °C.

***Nano-synbody formation.*** The 1-arm, 2-arm, and 3-arm nano-synbodies were prepared by mixing the respective 3-helix bundles (with single-stranded arms) in 1×PBS buffer (pH 7.4) with DNA-LCB1 conjugate at ratios of 1:1, 1:2, and 1:3, respectively. Subsequently, the mixture was incubated at 37 °C for 30 min and rapidly cooled down to 4 °C. After this, the assembled nanostructures were purified using a 50 kDa Amicon filter to remove any free DNA-LCB1 conjugate.

***Characterization of 3-helix bundle and nano-synbody.*** The samples were subjected to electrophoresis on an 8% native PAGE gel, which was prepared in 1×TAE-12.5 mM $MgCl_2$ buffer in-house. The gels were run at a constant voltage of 200 V at 10 °C for 1.5 h, with 1×TAE-12.5 mM $MgCl_2$ used as the running buffer. Subsequently, the gels were post-stained with ethidium bromide (EtBr) solution.

***Nano-synbody complexation with RBD and spike proteins.*** Samples of a specific 1-arm, 2-arm, or 3-arm nano-synbody (50 µL, 200 nM) in 1×TBS buffer (20 mM, pH 7.4) were incubated at 4 °C for 30 min with RBD protein at ratios of 1:1, 1:2, and 1:3, respectively. Similarly, the 3-arm nano-synbody sample (50 µL, 200 nM) in 1×TBS buffer (20 mM, pH 7.4) was incubated at 4 °C for 30 min, with spike proteins (WT and Omicron) at ratios of 1:0.5, 1:1, and 1:2.

***Characterization of nano-synbody complexation with RBD and spike proteins.*** The binding ability of nano-synbody with RBD and spike proteins was characterized using an electrophoretic mobility shift assay (EMSA). The samples were run on 1% agarose gel, which was prepared in 1×TAE-12.5 mM $MgCl_2$ buffer, and pre-stained with EtBr. The gel was run at a constant voltage of 75 at 4 °C for 1.5 h, with 1×TAE-12.5 mM $MgCl_2$ used as the running buffer.

### Toehold mediated strand displacement assay

DNA-LCB1 binding handles on the trivalent (3-arm) nano-synbody were extended at the 3′ end with three orthogonal 10-nt toehold domains. The 3-arm nano-synbody was assembled by thermal annealing and incubated with either wild-type (WT) or Omicron SARS-CoV-2 spike trimers at 37 °C for 30 min to form bound complexes, which were confirmed by native PAGE. Strand displacement reactions were performed by incubating the nano-synbody/spike complexes with complementary invader strands (S1, S2, and/or S3) at 37 °C for 10 min. Addition of one, two, or three invader strands displaced the corresponding number of DNA-LCB1 arms, yielding bi-, mono-, or zero-valent constructs, respectively. Strand-displaced constructs were analyzed by native gel-based binding assays. **Figure S4** shows the design and sequences of all strands for the toehold displacement assay.

### AFM imaging

The samples were adjusted to a concentration of 10 nM and deposited on freshly cleaved mica (Ted Pella, Inc. USA), followed by addition of 60 µL of 1×TAE buffer containing: 45 mM Tris pH 8.0, 12.5 mM $Mg(OAc)_2 \cdot 6H_2O$, and 2 mM EDTA, and then addition of 2 µL of 10 mM $NiCl_2$. The sample was allowed to adsorb at RT for 5 min, and then scanned in tapping mode on a Pico-Plus AFM (Molecular Imaging, Agilent Technologies) with NPS tips (Veeco, Inc. USA). All images were collected under ambient conditions in tapping mode.

### TEM imaging

5 µl of purified samples were applied on a commercially supplied formvar stabilized carbon type-B, 400mesh copper grids (Ted Pella, part number 01814-F) that had been glow discharged for 1 min at 15 mA using a Pelco easiGlow glow-discharge system (Ted Pella, Redding, CA, USA). Grids were stained using 5 µL of a freshly prepared 2% aqueous uranyl formate solution containing 25 mM sodium hydroxide (NaOH), and samples were incubated for 15 to 300 s depending on the concentration of the sample. Excess liquid was wicked away with Whatmann filter paper and grids left to dry for 30-60 min prior to imaging.

Images were acquired on a Talos microscope (Thermo Fisher LMT) operated at 120 kV accelerating voltage, using a charge-coupled device (CCD) camera at 73000x magnification.

Particles (see details in Table S3) were manually picked and class averaged using Relion3.0 software without CTF correction.

**Pseudovirus production and neutralization assay**

HIV-1 based SARS-CoV-2 pseudotyped virions were produced as described[1]. To produce the luciferase labeled pseudovirus, HEK293T cells were co-transfected with spike envelope plasmid (HDM-SARS2-spike-delta21, Addgene, Cat. 155130) or plasmids encoding a given spike variant (pcDNA3.3-SARS2-B.1.617.2, Addgene, Cat. 172320, pcDNA3.3_SARS2_omicron_BA.1 Addgene, Cat.180375, pcDNA3.3_SARS2_omicron BA.2 Addgene, Cat.183700, pCAGGS SARS-CoV-2 BA.4/5 Spike Addgene, Cat.18603), a packaging plasmid (psPAX2, Addgene, Cat. 12260), and a backbone plasmid (pLenti CMV-puro-luc, Addgene, Cat. 17477). All plasmids were transfected using Lipofectamine 3000. The medium was replaced with fresh medium after 12 h, and supernatants were harvested at 48 h post-transfection. In addition, a GFP report vector was used to replace the CMV-puro-luc to produce the GFP labeled pseudovirus. The lentivirus was concentrated by polyethylene glycol 8000 (PEG 8000) precipitation and the titer was determined by serial dilution transduction of hACE2-293T cells in white 96-well plates.

For the neutralization assay, pseudoviruses were incubated with serial dilutions of the nano-synbody for 30 min at 37 °C, and then added into the 96-well plate pre-seeded with $2\times10^4$ hACE2-293T cells per well. Cells without viruses or nano-synbody were used as blank controls, and cells with viruses but without nano-synbody were used as positive controls for virus infection. After 48 h incubation at 37 °C with 5% $CO_2$, cells were processed with luminescent regent (ONE-GloTM, Promega) according to the manufacturer's instructions, and luminescence (RLU) was measured with a microplate reader (Biotek Synergy H1). Inhibition was calculated using the following equation:

100- ([RLU of pseudovirus + nano-synbody]- [RLU of blank])/ ([RLU of pseudovirus only] – [RLU of blank])*100.

**Nanoparticle tracking analysis**

The purified pseudovirus sample was characterized for size distribution through nanoparticle tracking analysis (NTA), using a Nanosight NS300 system (Malvern Technologies, Malvern, UK). Briefly, the pseudovirus sample was diluted between 100- and 500-fold in freshly filtered (particle free) 1×PBS buffer (pH 7.4). This dilution ensured that concentration of sample remained within the range of $1 \times 10^8$-$10^9$ particles/mL. Subsequently, a volume (1 mL) of prepared sample was introduced into sample chamber and analysis was performed at a constant temperature of 25 °C. Videos for NTA analysis were recorded for 45 s. This process was replicated three times, maintaining a camera level of 14. The recordings were processed using NTA 3.4 software to determine the mean size and concentration of sample (**Figure S8B**).

**Gold surface preparation**

The bare glass coverslip was rinsed by acetone and DI water then dried by nitrogen flow. A 2-nm chromium layer and 47-nm gold layer were deposited by an E-beam thermal evaporator (PVD-75, Kurt J. Lesker) on a cleaned glass surface sequentially in a vacuum of $5 \times 10^{-6}$ Torr. The bare gold sensor chip was rinsed with ethyl alcohol and DI water, then dried by nitrogen and annealed using a hydrogen flame. The prepared gold sensor chips were placed in a standing jar for the following surface functionalization. A 5 mM 11-mercaptoundecanoic acid solution in pure ethyl alcohol was used to coat the carboxyl group on bare gold surface. The pH of the solution was adjusted using 37% HCl to ~2 to enhance the solubility of the 11-mercaptoundecanoic acid. To dispense the molecules better, the solution was sonicated for 10 min before introducing to the gold surface. The cleaned bare gold chips were immersed in this solution and sealed in the standing jar for 48 h to form the self-assemble monolayer (SAM) on gold surface. The carboxylated gold chips were rinsed with ethyl alcohol and DI water, then dried using a flow of nitrogen and stored under a nitrogen environment for future use.

#### SPR detection

The SPR experiments were performed on a home-built Kretschmann configuration setup. A 670-nm superluminescent diode (260-UHP-TOW2-PD, Superlum) was used to excite the SPR and the reflection light was detected by a CMOS camera (MC023MG-SY-UB, XIMEA) at 1 frame per second (FPS). The flow channel was pressed on the carboxylated gold surface, and the buffer flow was driven by a syringe pump (Fusion 200-X, Chemxy Inc.). An injection valve with isolated sample loop (C1CF-2346, VICI Inc.) was used to deliver the ligands.

The carboxylated gold surface was activated using NHS and EDC in order to immobilize proteins via their lysine residues. 3 mg NHS and 20 mg EDC were dissolved in 500 μL DI water and flowed over the carboxylated gold surface for 15 min at 5 $\mu L \cdot min^{-1}$. After removing the remaining NHS and EDC, 10 $\mu g \cdot mL^{-1}$ spike protein solution in 1×PBS was flowed onto the NHS activated gold surface for a 15 min immobilization at 5 $\mu L \cdot min^{-1}$, then the unbound NHS groups were terminated by 1 M ETA·HCl (pH 8.5) for 7 min at 10 $\mu L \cdot min^{-1}$. This surface with immobilized spike proteins was exposed to different ligands (antibodies and synbodies) and the intensity of the reflected light was recorded and averaged over the full frame to obtain the SPR response and determine the binding kinetics.

### S2. Simulations of nano-synbody with and without spike protein

#### All-atom molecular simulation of spike protein and LCB1 for B-factor acquisition

We performed all-atom simulations of two spike protein conformations and the LCB1 protein using Gromacs. The spike protein conformation, featuring two open (“up”) receptor binding domains (RBDs) and one closed (“down”) RBD, was obtained from PDB ID: 7BNO. The conformation with three open RBDs was sourced from PDB ID: 7UHC. The LCB1 protein was extracted from PDB ID: 7JZM. The highly flexible residues absent in the spike crystal structures were modeled using the CHARMM-GUI PDB reader. We employed the CHARMM27 forcefield and tip3p water model. After solvation, salt ions were introduced to neutralize the charge in the box. The structures underwent relaxation via steepest descent minimization, followed by a 1-ns NVT equilibration at 300 K, and a 1-ns NPT equilibration at a reference pressure of 1 bar. Production simulations spanned 100 ns. The trajectories were centered, and periodic boundary effects removed. The RMSF and RMSD values were aligned with, and calculated relative to, the backbone of the mean structure, excluding the initial 10 ns. The conformation with the smallest RMSD value was selected for coarse-graining.

#### Protein coarse-graining and equilibration

The spike and LCB1 proteins underwent coarse-graining using the ANMUtils module within the ANM-oxDNA software package. This process generated structure and parameter files in an oxDNA-compatible format. The parameter file, derived from the B-factor/RMSF data obtained from the all-atom simulations, was used to parameterize an anisotropic network model that governs the dynamics of the proteins. The coarse-grained proteins were relaxed though ANM-oxDNA molecular dynamics simulations using the ipy-oxDNA package. The simulations were conducted with a time-step of 0.606 femtoseconds (fs), a temperature of 293.15 K, a salt concentration of 1 M, a max backbone force of 486.3 pN, and a diffusion coefficient of 2.5, under an Anderson-like thermostat for 1 x $10^5$ steps. Following relaxation, the proteins were equilibrated under the same conditions, with the exception of a time-step of 6.06 fs and no limit to the backbone force. This equilibration phase lasted for 5 x $10^6$ steps.

**Nano-synbody coarse-graining and equilibration**

We utilized the TacoxDNA extension in oxView to convert the Tiamat2 nano-synbody designs into the oxDNA format. The structure was relaxed interactively in oxView, initially applying a harmonic trap external potential between all native base pairs with a stiffness of 171.27 pN/nm. Subsequently, a Monte Carlo simulation with a max backbone force of 486.3 pN was conducted until the backbone distances were adequately relaxed, spanning approximately 1 x $10^5$ steps. Monte Carlo simulations without a backbone force limit were then executed for about 5 x $10^5$ steps. The nano-synbody underwent equilibration, after the base pair external potentials were removed, using molecular dynamics simulations performed with the ANM-oxDNA model for approximately 1 x $10^6$ steps. These simulations were conducted with a time step of 6.06 fs, a temperature of 293.15 K, and a salt concentration of 1 M. To maintain a constant temperature throughout the simulations we applied an Anderson-like thermostat. A diffusion coefficient of 2.5 simulation units was used. We note that there is not a direct correspondence between coarse-grained model simulation time and actual time in experiment. Prior work has shown that the timescale of oxDNA is about 100 times faster than the corresponding time in seconds when converted from the model units.[2]

**Coarse-grained modeling of DNA-protein and protein-protein interactions**

The DNA-LCB1 conjugate linkage—via the sulfosuccinimidyl 4-(N-maleimidomethyl) cyclohexane-1-carboxylate (SMCC) bifunctional crosslinker—was modeled using a skewed Gaussian potential between the C-terminal amino acid and the 3' nucleotides. The potential was parameterized by simulating the linker in all-atom molecular dynamics. The simulation used Gromacs with the opls-as force field with 1.14*CM1A-LBCC partial charges using the LigParGen web server to generate the input. The simulations were run at 300 K for 50 ns across three replicas, following NVT and NPT equilibration. The solvent used was SPC/E water. We measured the end-to-end distance of the linker over the simulations at 0.5-ns intervals across all three simulations. We then matched this end-to-end distance distribution in oxDNA by sampling two particles attached by a skewed Gaussian potential over a range of skew parameters and used the best match. The bound state of the LCB1 with respect to the RBD region on the spike protein was modeled using two external harmonic potentials. These potentials were applied between the closest LCB1 and RBD contact residues within the 7JZM crystal structure, thereby enforcing no rotation.

**Simulation of nano-synbody bound to the spike protein**

The coarse-grained system with the spike protein and three LCB1 proteins was created in oxView via drag-and-drop. The LCB1 proteins were positioned into the RBD binding pockets, and harmonic traps were applied. The system was saved in oxView, generating a new parameter file that accounted for all the proteins. The system was then relaxed and equilibrated in a manner similar to the individual proteins. Next, the nano-synbody was dropped into oxView, positioned next to the LCB1 proteins, and harmonic traps were applied between the C-terminal and 3' nucleotide. The system was relaxed as previously described. The harmonic trap between the LCB1 proteins and DNA was then replaced with a skewed gaussian potential. The system was equilibrated for 1 x $10^7$ steps with a time step of 6.06 fs. Six replicas were run for each of the three systems: the nano-synbody with three LCB1 proteins bound to the two RBD open spike, bound to the three RBD open spike, and unbound. Each replica was run for 3 x $10^9$ steps.

#### LCB1 center of mass distance histograms

We applied three observables to each system to calculate the distance between the centers of mass of the LCB1 proteins with respect to each other. The center of mass distance observables were saved every 1 x $10^3$ steps. The distance data of each replica was binned into a histogram with 200 equally spaced bins for all systems. When plotted, the probability of a bin was given by the mean probability across the six replicas, with the error bars represented by the standard deviation of the mean. To calculate the mean distance for each system, we first computed the mean for each replica. We then calculate the average of these means, which we took as the overall average distance. The standard deviation of these means was used as the standard error of the mean.

#### Molecular visualization

We employed oxDNA Analysis Tools to calculate the mean structure, the root mean-squared fluctuations (RMSF), and the root mean-squared deviation (RMSD). The mean oxDNA structures were converted into PDB using tacoxDNA. For the electrostatic surface rendering and RMSF visualization of the structures, we utilized ChimeraX.

**All simulation files are available on:** https://github.com/sulcgroup/nanosynbody

### S3. Design and DNA sequences used for nano-synbody synthesis.

**A**

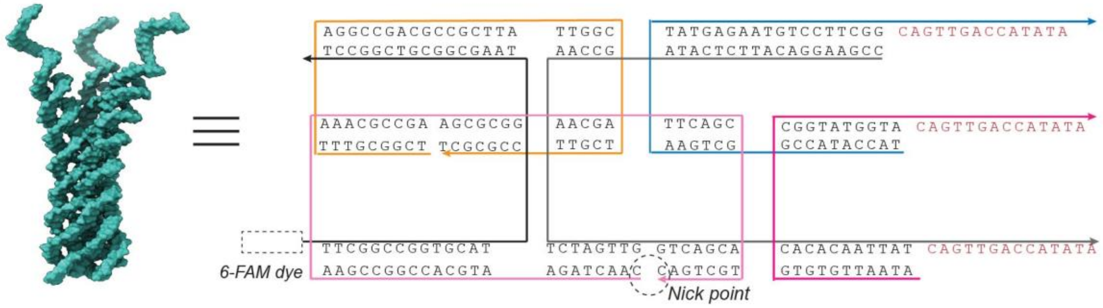


**B**

| Name | Sequence (5'→3') |
|---|---|
| Organge | 5'-TCG GCG TTT AGG CCG ACG CCG CTT ATT GGC TGC TTC CGC GCT-3' |
| Black | 5'-TTC GGC CGG TGC ATT AAG CGG CGT CGG CCT-3' |
| Ash | 5'-CCG AAG GAC ATT CTC ATA GCC AAT CTA GTT GGT CAG CAC ACA CAA TTA TCA GTT GAC CAT ATA-3' |
| Blue | 5'-TAC CAT ACC GGC TGA ATA TGA GAA TGT CCT TCG GCA GTT GAC CAT ATA-3' |
| Pink | 5'-CAA CTA GAA TGC ACC GGC CGA ATT AAA CGC CGA AGC GCG GAA GCA TTC AGC TGC TGA C-3' |
| Fuchsia | 5'-ATA ATT GTG TGC GGT ATG GTA CAG TTG ACC ATA TA-3' |

**C**

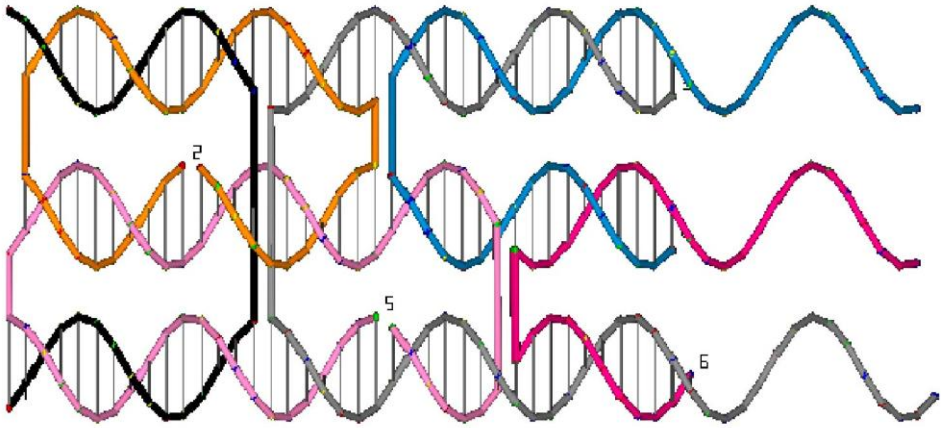

**Figure S1. Strand routing diagram and sequences used for the 3-arm nano-synbody. A)** Representative schematic and strand routing diagram for the 3-arm nano-synbody. **B)** Sequences of ssDNA strands used, wherein the green box region indicates the complementary sequences to the DNA-LCB1 conjugate. **C)** Tiamat design of 3-arm nano-synbody, showing the helicity and routing paths of the six strands. The colors correspond to the strands as indicated in (**A**).

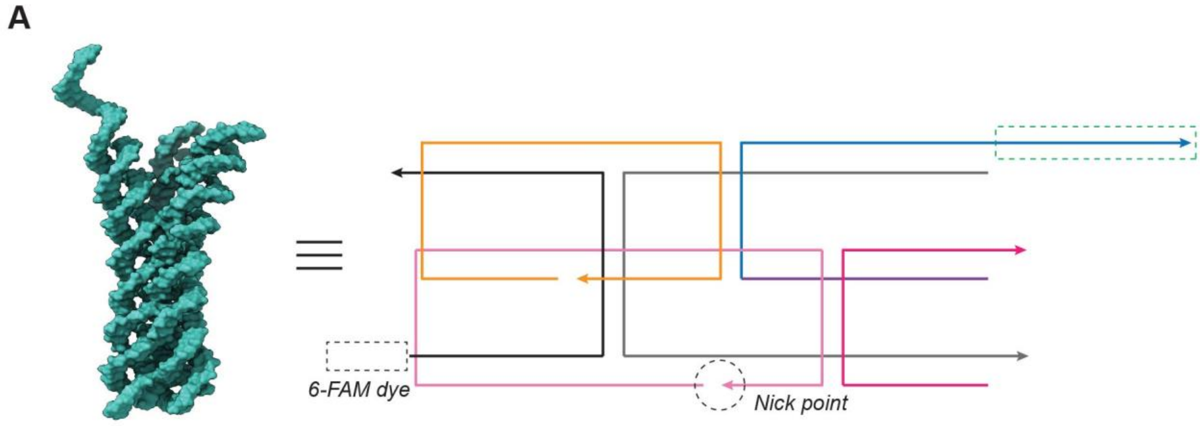


**B**

| Name | Sequence (5'→3') |
|---|---|
| Organge | TCG GCG TTT AGG CCG ACG CCG CTT ATT GGC TGC TTC CGC GCT |
| Black | 6-FAM-TTC GGC CGG TGC ATT AAG CGG CGT CGG CCT |
| Ash | CCG AAG GAC ATT CTC ATA GCC AAT CTA GTT GGT CAG CAC ACA CAA TTA TTT T |
| Blue | TAC CAT ACC GGC TGA ATA TGA GAA TGT CCT TCG GCA GTT GAC CAT ATA |
| Pink | CAA CTA GAA TGC ACC GGC CGA ATT AAA CGC CGA AGC GCG GAA GCA TTC AGC TGC TGA C |
| Fuchsia | ATA ATT GTG TGC GGT ATG GTA TTT |

**Figure S2. Strand routing diagram and sequences used for the 1-arm nano-synbody. A)** Representative schematic and strand routing diagram for the 1-arm nano-synbody. **B)** Sequences of ssDNA strands used, wherein the green box region indicates the complementary sequences to the DNA-LCB1 conjugate.

A

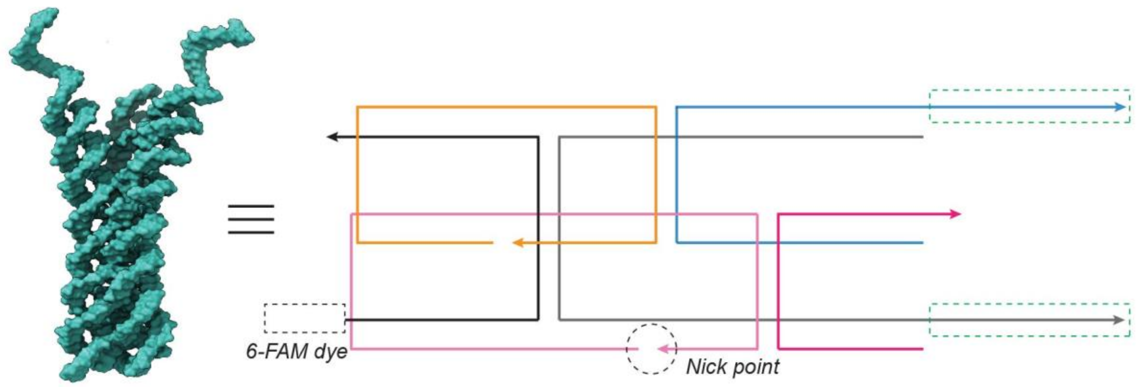


B

| Name | Sequence (5'→3') |
|---|---|
| Organge | TCG GCG TTT AGG CCG ACG CCG CTT ATT GGC TGC TTC CGC GCT |
| Black | 6-FAM- TTC GGC CGG TGC ATT AAG CGG CGT CGG CCT |
| Ash | CCG AAG GAC ATT CTC ATA GCC AAT CTA GTT GGT CAG CAC ACA CAA TTA TCA GTT GAC CAT ATA |
| Blue | TAC CAT ACC GGC TGA ATA TGA GAA TGT CCT TCG GCA GTT GAC CAT ATA |
| Pink | CAA CTA GAA TGC ACC GGC CGA ATT AAA CGC CGA AGC GCG GAA GCA TTC AGC TGC TGA C |
| Fuchsia | ATA ATT GTG TGC GGT ATG GTA TTT |

**Figure S3. Strand routing diagram and sequences used for the 2-arm nano-synbody. A)** Representative schematic and strand routing diagram for the 2-arm nano-synbody. **B)** Sequences of ssDNA strands used, wherein the green box region indicates the complementary sequences to the DNA-LCB1 conjugate.

**A**

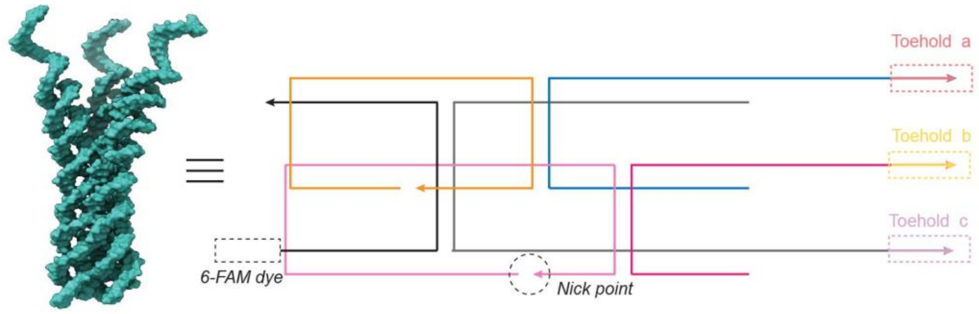


**B**

| Name | Sequence (5'→3') |
|---|---|
| Organge | TCG GCG TTT AGG CCG ACG CCG CTT ATT GGC TGC TTC CGC GCT |
| Black | 6-FAM-TTC GGC CGG TGC ATT AAG CGG CGT CGG CCT |
| Ash | CCG AAG GAC ATT CTC ATA GCC AAT CTA GTT GGT CAG CAC ACA CAA TTA TCA GTT GAC CAT ATA CTT GTC TCGA |
| Blue | TAC CAT ACC GGC TGA ATA TGA GAA TGT CCT TCG GCA GTT GAC CAT ATA AGT TCC AGGA |
| Pink | CAA CTA GAA TGC ACC GGC CGA ATT AAA CGC CGA AGC GCG GAA GCA TTC AGC TGC TGA C |
| Fuchsia | ATA ATT GTG TGC GGT ATG GTA CAG TTG ACC ATA TA AAA CAG GGCT |
| Invader S1 | TCG AGA CAA GTA TAT GGT CAA CTG |
| Invader S2 | TCC TGG AAC TTA TAT GGT CAA CTG |
| Invader S3 | AGC CCT GTT TTA TAT GGT CAA CTG |

**Figure S4. Reversible protein binding via toehold-mediated strand displacement. A)** Schematic strand routing diagram for the trivalent (3-arm) nano-synbody incorporating orthogonal 3′ toehold domains for programmable strand displacement. **B)** Sequences of ssDNA strands used in the assay; boxed regions denote the three orthogonal toehold domains that initiate strand displacement.

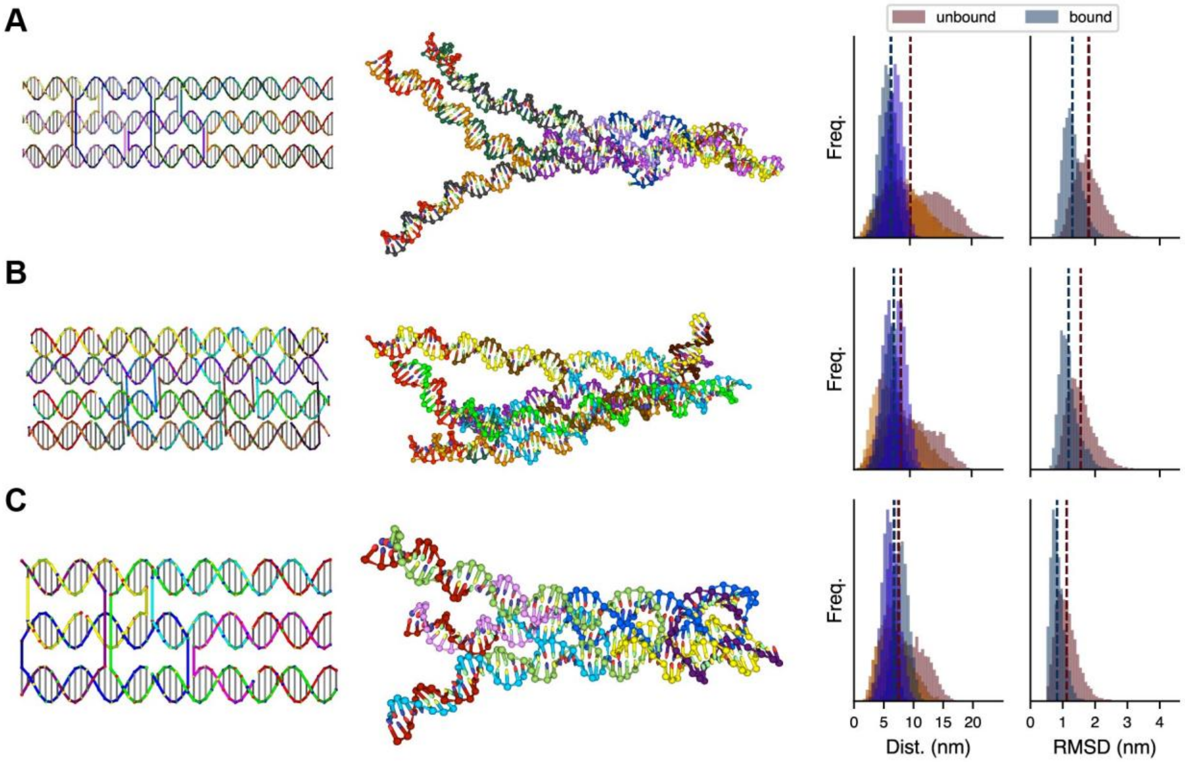


**Figure S5. Simulation-guided assessment of three potential DNA nano-synbody designs. A)** 3-helix bundle with arms emerging from a double-crossover tile motif. **B)** 4-helix bundle based on a honeycomb lattice cross-section, with three of the arms containing conjugation sites for the binding protein. **C)** 3-helix bundle with shorter arms and a truncated body to increase the rigidity of the structure. In each of the Tiamat designs (left), the strands for protein attachment are labeled in red. The designs were converted to oxDNA format, relaxed, and simulated both independently (representative configuration shown center) and bound to the WT spike protein. The quality of the design was assessed by comparing the distance between the functionalized nucleotides and the overall RMSD from the average structure between the unbound and bound states (right). The shorter 3-helix bundle (**C**) was chosen due to its overall higher rigidity and better match between bound and unbound inter-arm distances.

### S4. Additional characterization data.

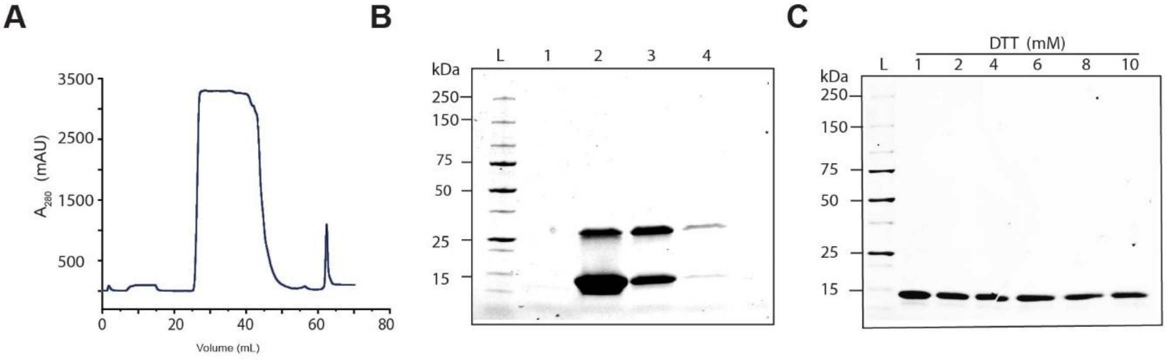


**Figure S6. Purification and characterization of LCB1**. **A)** Elution trace of LCB1 using fast protein liquid chromatography (FPLC), monitored by in-line absorption at 280 nm. The peak from 60-65 mL was collected as eluted fractions (at 1 mL per fraction). **B)** SDS-PAGE gel analysis of the collected fractions; the LCB1 protein appears as a disulfide-linked dimer band (*upper*) and a monomer band (*lower*). **C)** SDS-PAGE gel analysis of the disulfide-reduced LCB1, by increments of DTT concentration (lanes (*L*-*R*): ladder, 1, 2, 4, 6, 8, 10 mM DTT).

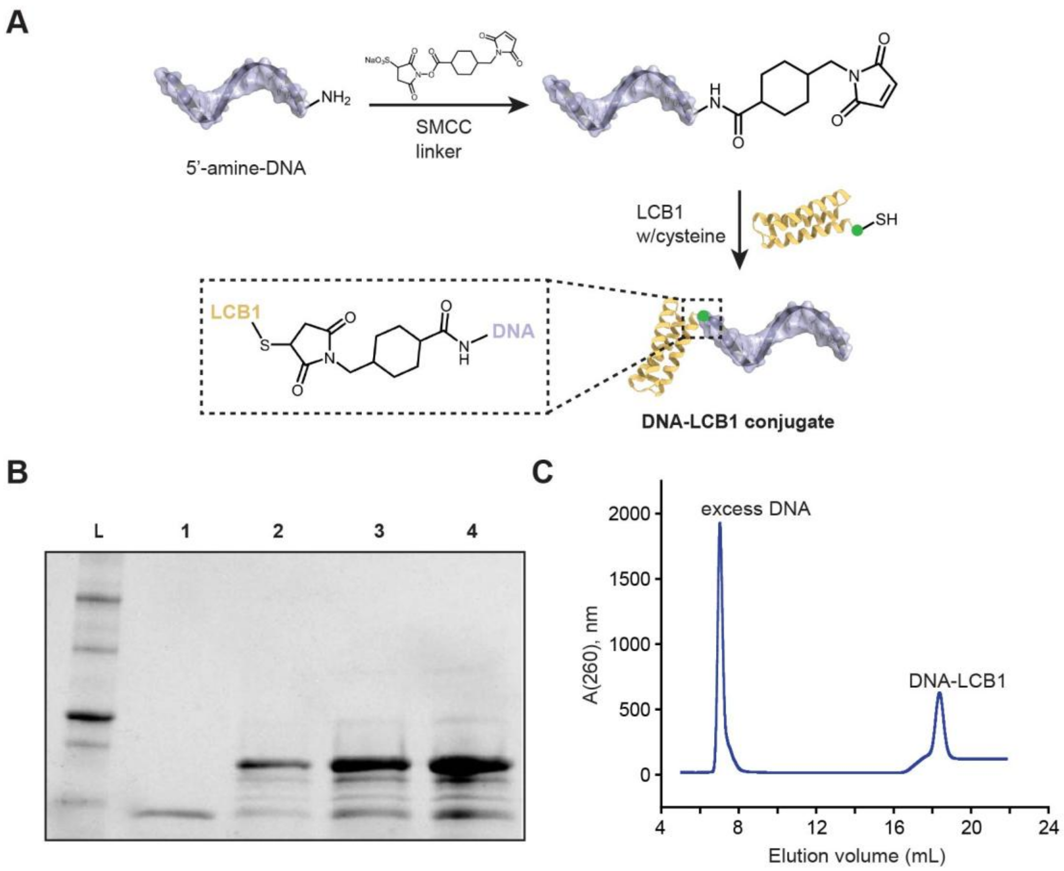


**Figure S7. DNA-LCB1 conjugation and purification. A)** Schematic representation depicting the site-specific functionalization strategy of LCB1 (*yellow*) with a DNA strand (*light purple*). LCB1 is modified and expressed with a cysteine residue (*green*) conjugated to a DNA functionalized with a maleimide via the SMCC crosslinker. **B)** SDS-PAGE gel analysis of the DNA-LCB1 mixture with varying fold molar excess of maleimide-functionalized ODN. Lanes: L, ladder; 1, LCB1; 2, LCB1:DNA = 1:1; 3, LCB1: DNA = 1:2; 4, LCB1: DNA = 1:5. **C)** Purification of the DNA-LCB1 conjugate via fast protein liquid chromatography (FPLC) system equipped with a His-trap column. The elution profile shows the first peak corresponding to excess free DNA, while the second peak represents the DNA-LCB1 conjugate that was collected and further analyzed.

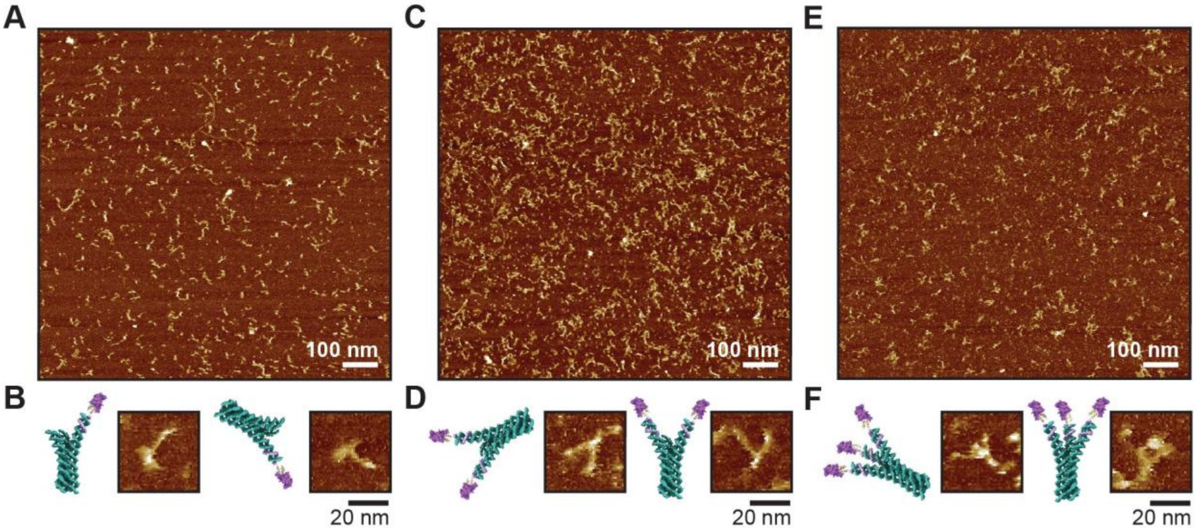


**Figure S8. AFM characterization of nano-synbody structures. A)** Overview AFM image of the 1-arm nano-synbody. **B)** Schematic representations depicting distinct views of the 1-arm nano-synbody, alongside their respective zoomed-in AFM images. **C)** Overview AFM images of the 2-arm nano-synbody. **D)** Schematic representations depicting distinct views of the 2-arm nano-synbody, alongside their respective zoomed-in AFM images. **E)** Overview AFM images of the 3-arm nano-synbody. **F)** Schematic representations depicting distinct views of the 3-arm nano-synbody, alongside their respective zoomed-in AFM images at higher magnification. For enhanced contrast on AFM, all samples were incubated with the RBD protein (*purple*) prior to imaging.

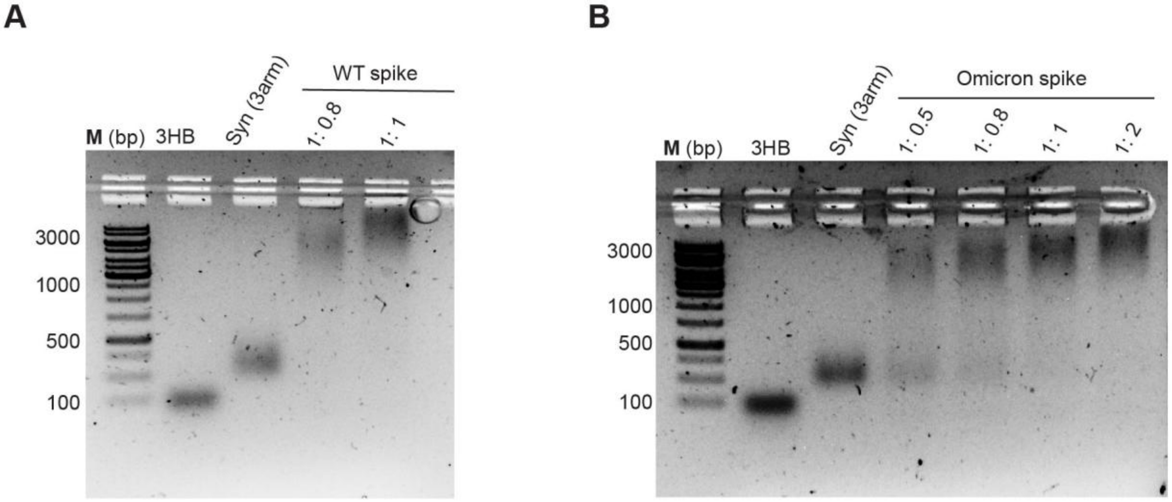


**Figure S9. Evaluation of 3-arm nano-synbody binding to WT and Omicron spike proteins via electrophoretic mobility shift assays (EMSA) on a 1% agarose gel.** The 3-arm nano-synbody was co-incubated with either WT or Omicron spike at 37 °C in varying ratios (as labeled above the wells), with band shifts serving as an indicator of binding. **A)** EMSA depicting the binding between 3-arm nano-synbody and WT spike protein. **B)** EMSA illustrating the binding of 3-arm nano-synbody with Omicron BA.1 spike protein.

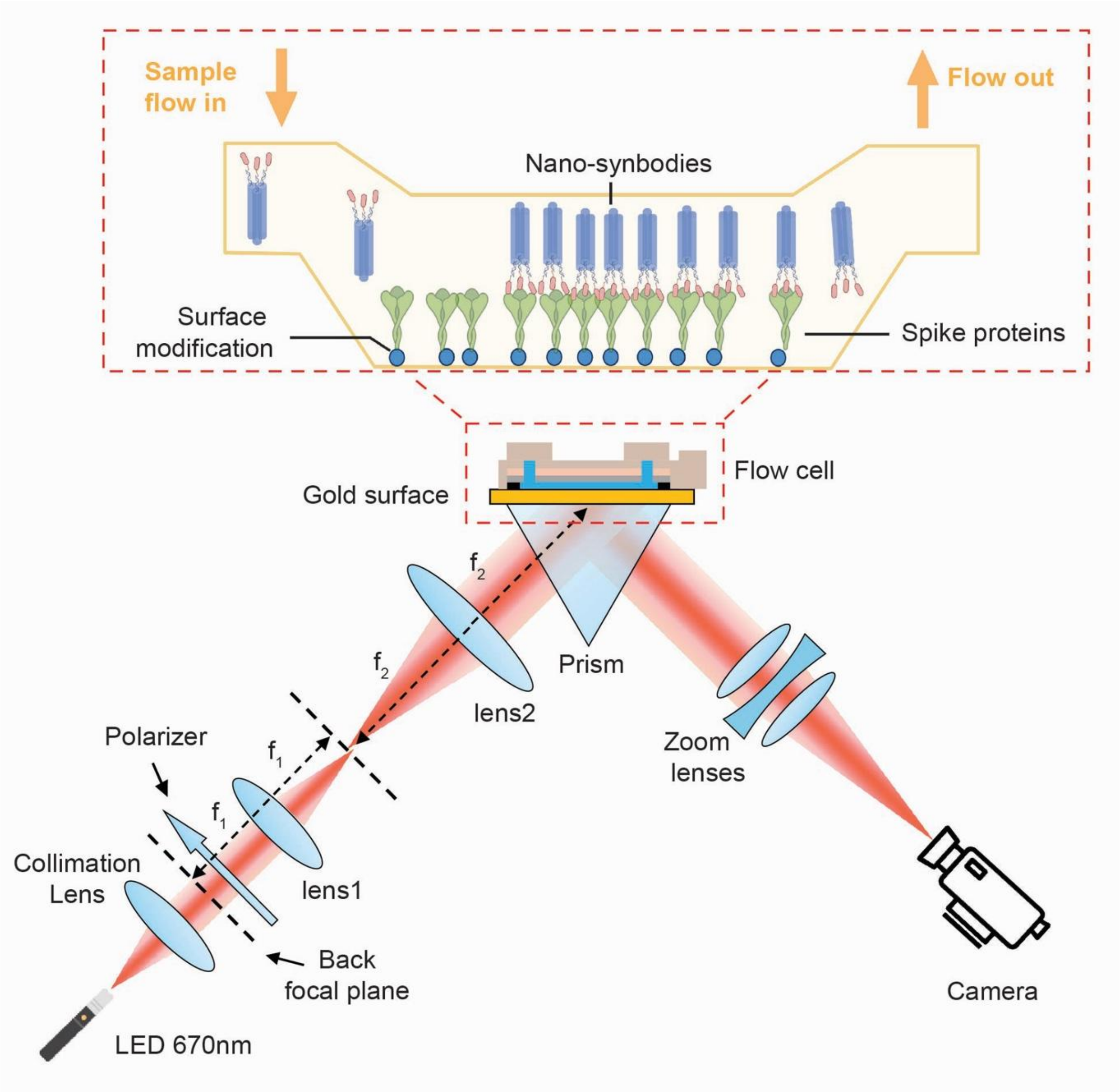


**Figure S10. Schematic illustration of the SPR setup**. Spike proteins are immobilized on a gold-coated sensor surface within a flow channel, and nano-synbodies are introduced under continuous flow to bind the surface-tethered proteins. Prism-coupled incident light excites surface plasmons at the metal–dielectric interface. Binding-induced refractive index changes shift the SPR condition, which is detected as changes in reflected light intensity and recorded in real time to determine association and dissociation kinetics.

| | WT spike | | | Omicron Spike | | |
|---|---|---|---|---|---|---|
| | $k_a$ (×$10^5$ $M^{-1}s^{-1}$) | k_d (×$10^{-5}$ $s^{-1}$) | K_D (×$10^{-12}$ M) | $k_a$ (×$10^5$ $M^{-1}s^{-1}$) | k_d (×$10^{-5}$ $s^{-1}$) | K_D (×$10^{-12}$ M) |
| **LCB** | 4.10 | 67.10 | 1640 | NA | NA | NA |
| **DNA-LCB** | 3.40 | 41.5 | 1220 | NA | NA | NA |
| **1arm** | 1.61 | 19.06 | 1180 | NA | NA | >100000 |
| **2arm** | 19.8 | 38.6 | 195 | 0.622 | 24.4 | 3920 |
| **3arm** | 62.0 | 6.89 | 11.2 | 22.4 | 21.4 | 95 |

**Table S1. SPR kinetics of synbody-spike interactions.** Association ($k_a$), dissociation ($k_d$), and equilibrium dissociation ($K_D$) constants for WT and Omicron spike proteins measured by SPR.

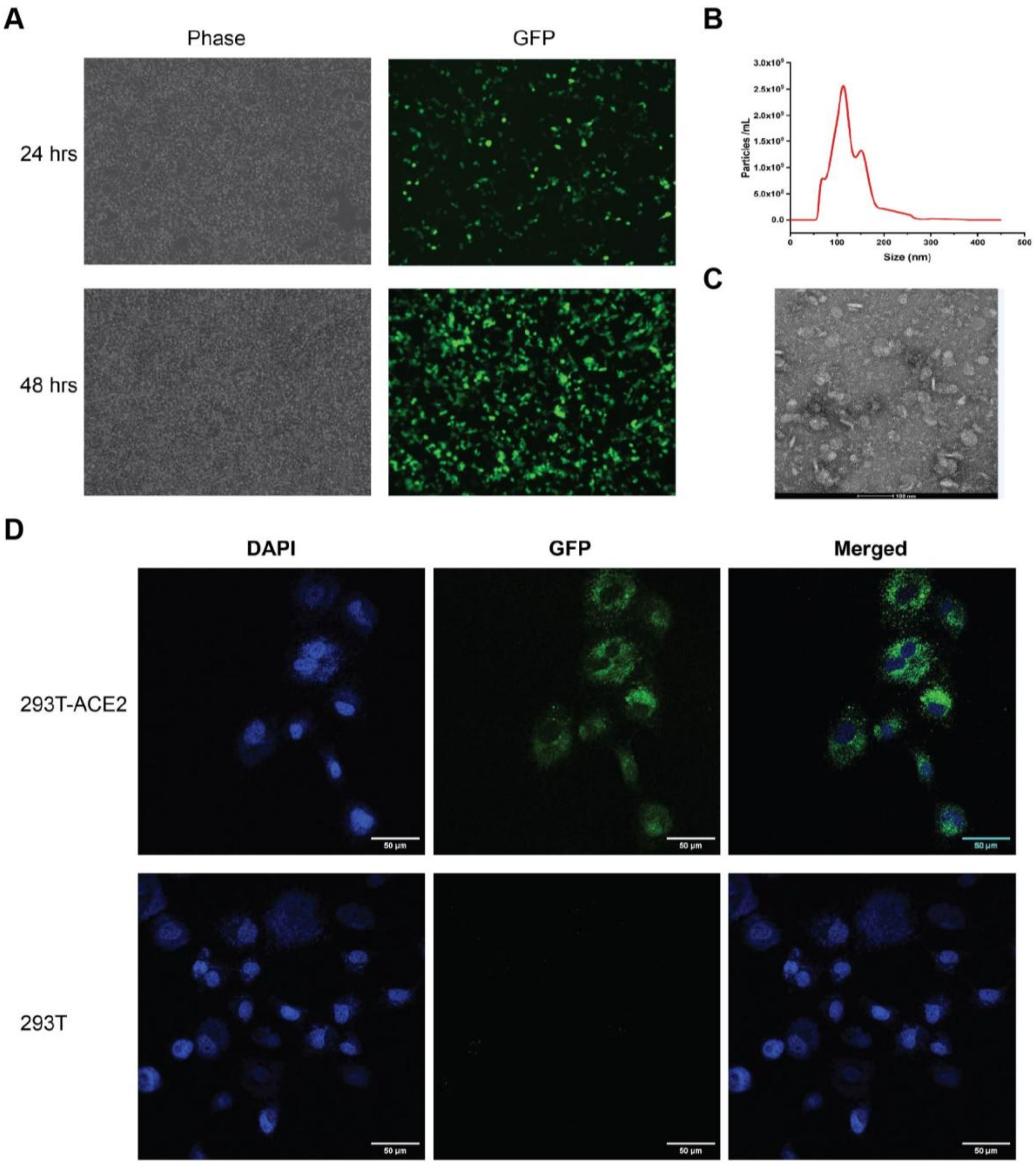


**Figure S11. Characterization and verification of SARS-CoV-2 spike pseudoviruses**. HEK293T cells were employed for the generation of SARS-CoV-2 spike pseudoviruses by means of transfection with plasmids encoding for the spike proteins along with the necessary packaging plasmids. The efficiency of this transduction process was subsequently evaluated via fluorescence microscopy for the delivered cargo (the GFP gene). **A)** Representative fluorescence microscopy data, visualizing the production of GFP-labelled SARS-CoV-2 pseudoviruses at 24 and 48 h. **B)** Nanoparticle tracking analysis data of the pseudoviruses post-PEG purification, indicating a mean size at ~100 nm. **C)** Negative-stain (ns) Transmission Electron Microscopy (TEM) imaging of the purified pseudoviruses, demonstrating the expected morphology previously reported. **D)** Fluorescence confocal images depict the infection of the 293T-ACE2 cell line and the 293T cell line by Pseudovirus. Scale bar: 50 μm.

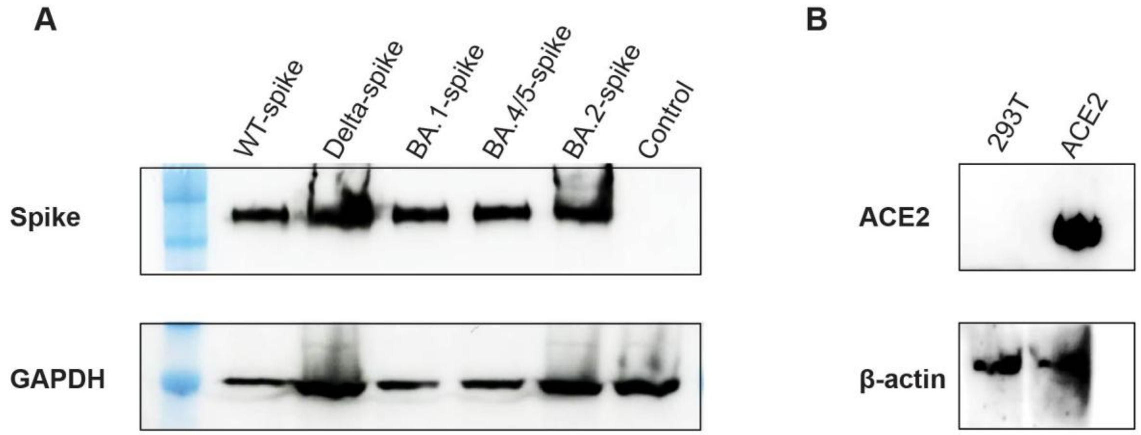


**Figure S12**. **Analysis of SARS-CoV-2 spike plasmid expression, and construction of an hACE2-293T stable cell line**. **A)** Diverse spike plasmids—including wild type (WT), Delta variant, and Omicron BA.1, BA.2, and BA.4/5 variants—were transfected separately into the 293T cell line; a negative control plasmid was also included. The resultant spike protein expression was probed via Western blotting. **B)** Western blotting was used to discern the difference in ACE2 protein expression between the original 293T cell line and the engineered hACE2-293T cell line.

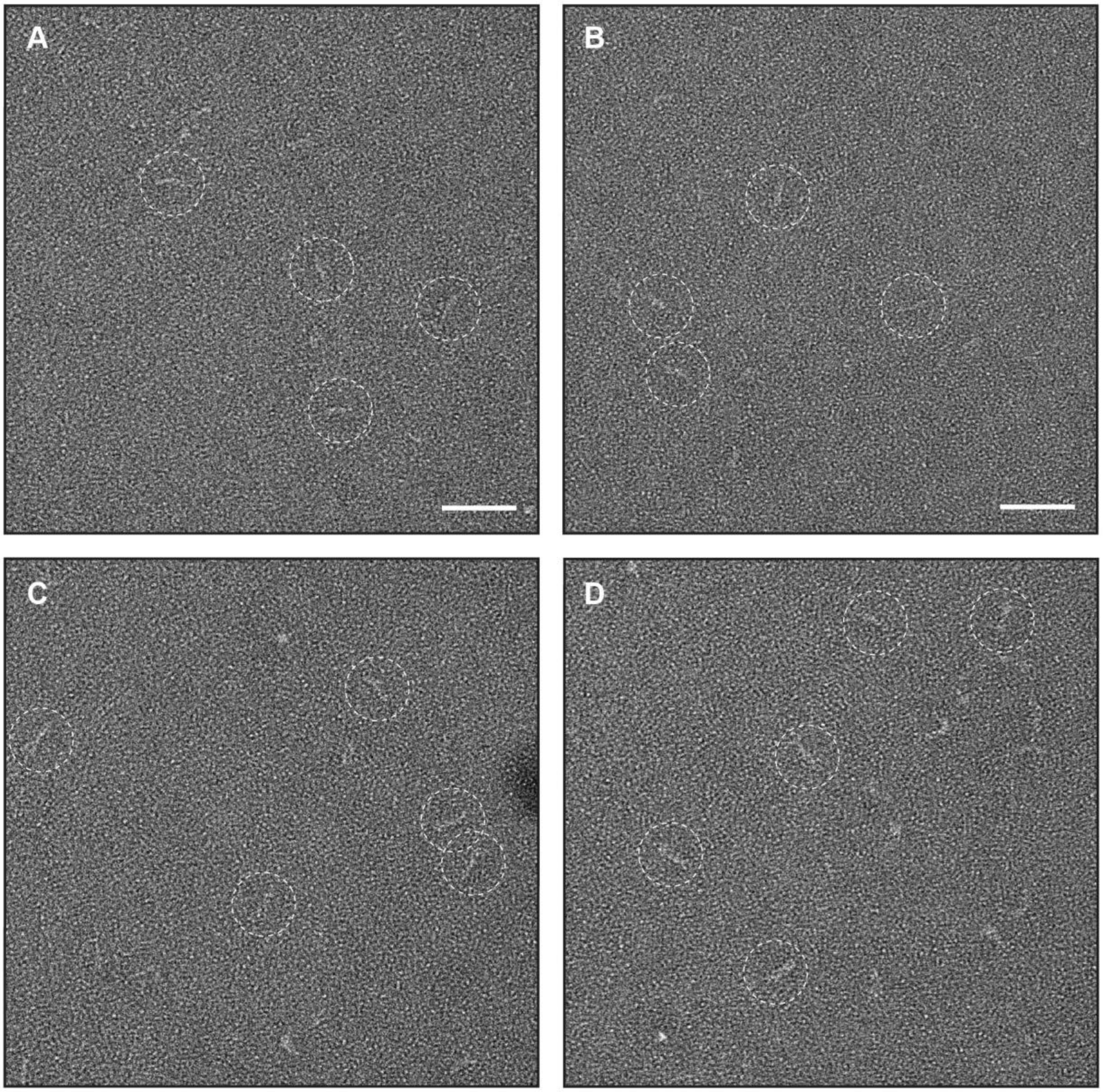


**Figure S13. Additional nsTEM images of 3-helix bundle and 3-arm nano-synbody. A-B)** Images of negatively-stained 3-helix bundle (3-arm), without LCB1 proteins attached, show well dispersed particles. White circles highlight representative particles. **C-D)** Images of negatively-stained well dispersed 3-arm nano-synbody (with LCB1 proteins attached). White circles highlight representative particles.

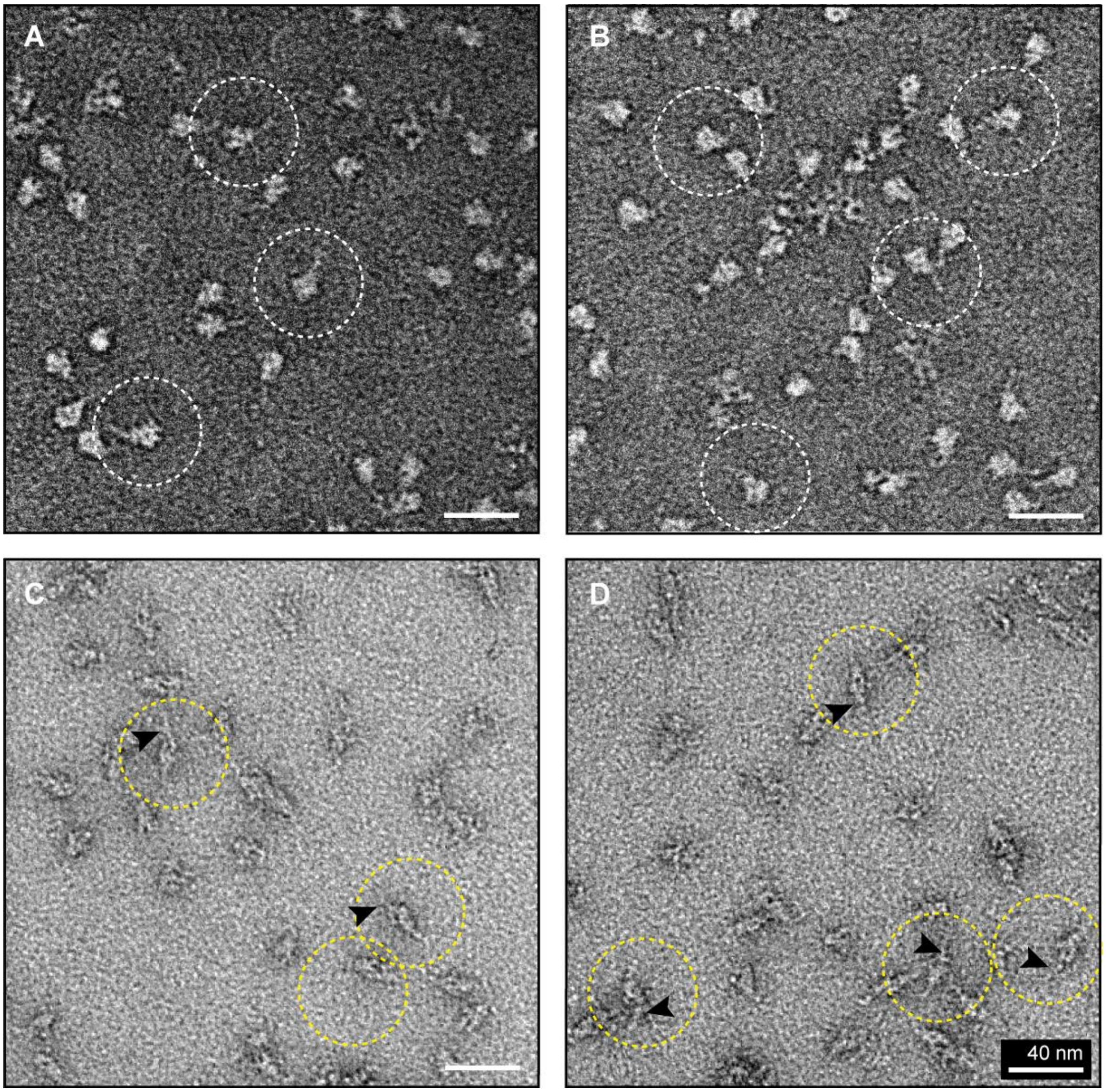


**Figure S14. nsTEM images of full-length WT spike protein and 3-arm nano-synbody/WT spike protein complex. A-B)** Images of negatively-stained spike particles. White circles highlight representative particles. **C-D)** Images of negatively-stained, well dispersed nano-synbody/WT spike protein complexes. Yellow circles highlight representative particles. Arrow indicates nano-synbody bound with the spike protein

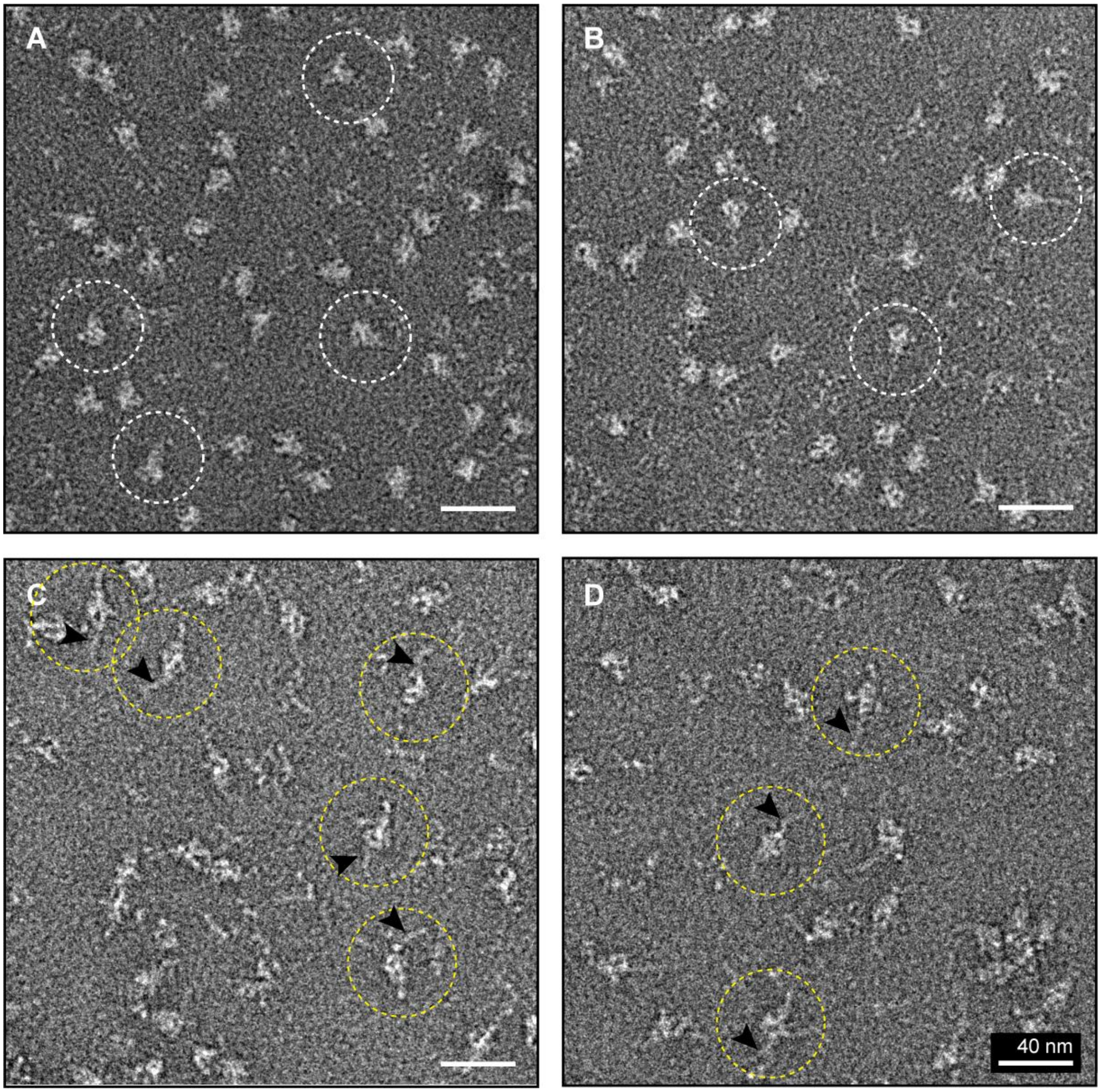


**Figure S15. nsTEM images of Omicron spike protein and 3-arm nano-synbody/Omicron BA.1 spike protein complex. A-B)** Images of negatively-stained spike particles. White circles highlight representative particles. **C-D)** Images of negatively-stained, well dispersed nano-synbody/Omicron BA.1 spike protein complexes. Yellow circles highlight representative particles. Arrow indicates nano-synbody bound with the spike protein

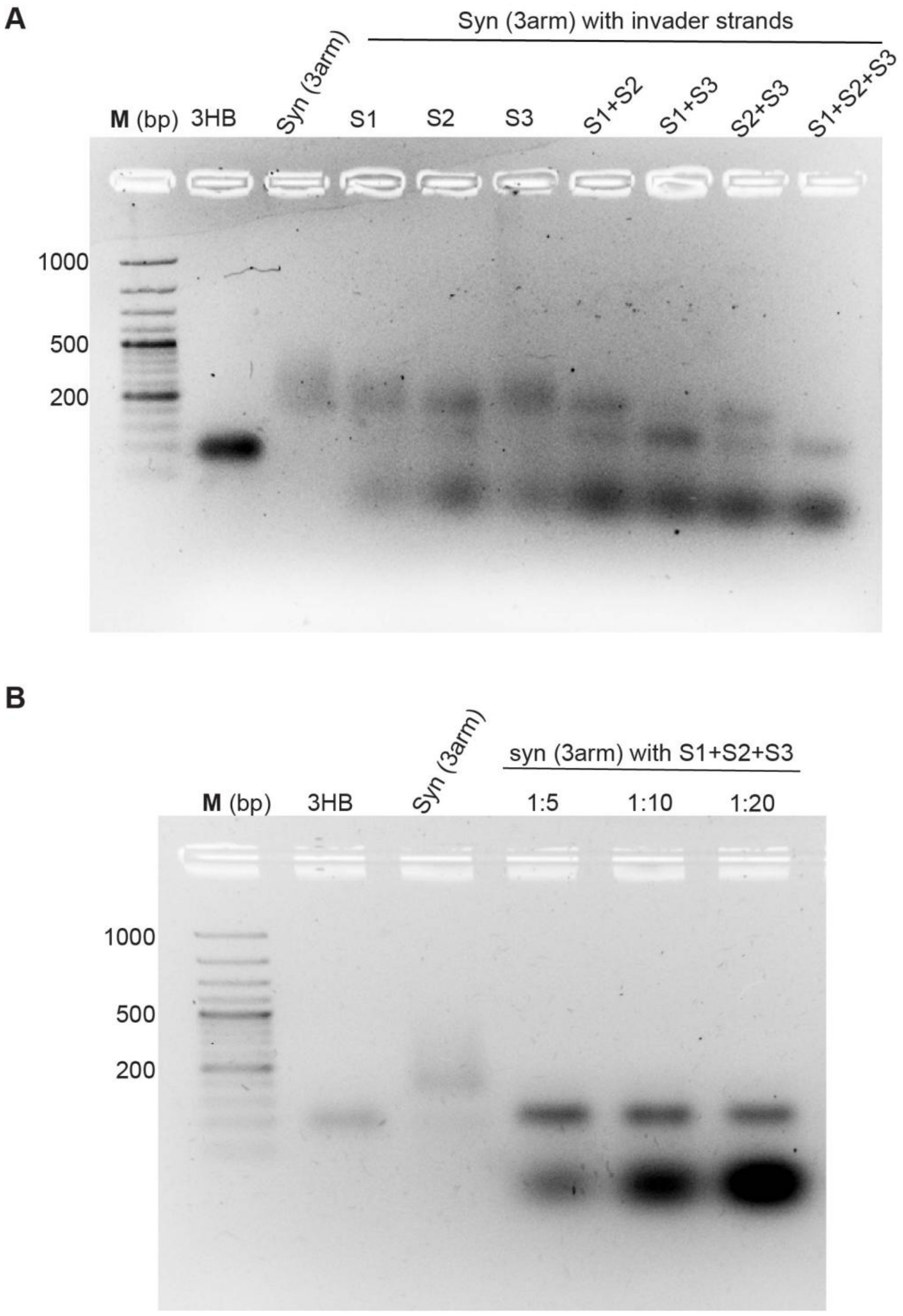


**Figure S16. Toehold-mediated strand displacement of the 3-arm nano-synbody. A)** Native PAGE analysis of the trivalent nano-synbody incubated with individual (S1, S2, S3), paired (S1+S2, S1+S3, S2+S3), or all three (S1+S2+S3) invader strands, showing stepwise displacement of DNA-LCB1 arms. **B)** Stoichiometry optimization for complete displacement using S1+S2+S3 at increasing nano-synbody: invader molar ratios (1:5, 1:10, 1:20).
